# LipoGlo: A sensitive and specific reporter of atherogenic lipoproteins

**DOI:** 10.1101/522615

**Authors:** James H. Thierer, Stephen C. Ekker, Steven A. Farber

**Affiliations:** Carnegie Institution for Science Department of Embryology, Baltimore, MD 21218; Johns Hopkins University Department of Biology, Baltimore, MD 21218; Mayo Clinic Department of Biochemistry and Molecular Biology, Rochester, MN 55905

**Keywords:** Lipoprotein, Apolipoprotein-B, ApoB, atherosclerosis, cardiovascular disease, zebrafish, pla2g12b

## Abstract

Apolipoprotein-B (APOB) is the structural component of atherogenic lipoproteins, lipid-rich particles that drive atherosclerosis by accumulating in the vascular wall. As atherosclerotic cardiovascular disease is the leading cause of death worldwide, there is an urgent need to develop new strategies to prevent lipoproteins from causing vascular damage. Here we report the LipoGlo system, which uses a luciferase enzyme (NanoLuc) fused to ApoB to monitor several key determinants of lipoprotein atherogenicity including particle abundance, size, and localization. Using LipoGlo, we are able to comprehensively characterize the lipoprotein profile of individual larval zebrafish and collect the first images of atherogenic lipoprotein localization in an intact organism. We discover multiple unexpected extravascular lipoprotein localization patterns, as well as identify *pla2g12b* as a potent regulator of lipoprotein size. ApoB-fusion proteins thus represent a uniquely sensitive and specific approach to study atherogenic lipoproteins and their genetic and small molecule modifiers.

## INTRODUCTION

ApoB-containing lipoproteins (ABCLs) are the etiological agents of atherosclerotic cardiovascular disease [1], which is the leading cause of mortality worldwide [2]. ABCLs serve to shuttle lipids throughout the circulation, but occasionally cross the vascular endothelium to form lipid-rich deposits within the vascular wall that develop into atherosclerotic plaques [1]. ABCLs are frequently characterized indirectly through measurement of their triglyceride and cholesterol content, and high-risk individuals with elevated lipid levels are prescribed lipid-lowering therapies such as statins [3]. Such drugs effectively reduce cardiovascular disease risk by lowering the levels of cholesterol carried by atherogenic lipoproteins (often called “bad cholesterol”).

Indirect (lipid-focused) measurements, however, provide very limited information on ABCL properties such as particle concentration or size distribution, both of which are key determinants of atherogenic potential. For example, serum Apolipoprotein-B (ApoB) levels directly reflect the concentration of ABCL particles and show a stronger correlation with cardiovascular disease risk than lipid metrics (including cholesterol) [4, 5]. The size distribution of lipoprotein particles is also relevant to cardiovascular disease risk, as there are numerous classes of ABCLs that can be differentiated by size and show varying degrees of atherogenicity [6]. Low-density lipoproteins (LDL) are the smallest and most abundant class of ABCLs and are thought to be the primary drivers of atherosclerosis. There is significant size variation within the LDL particle class, and smaller particles are associated with increased atherogenicity [7]. For example, approximately 25% of the adult population produces unnaturally small LDL particles, and as a result have ~3-fold higher risk for cardiovascular disease [8].

Many of the genetic and environmental factors governing ABCL size and abundance remain undiscovered or poorly characterized [9-11], and even fewer have been successfully targeted pharmaceutically [12-14]. It has proven particularly difficult to identify drugs that modulate ABCL size and abundance because the simplified model systems (such as cultured cells or invertebrate models) typically used in high-throughput drug screening do not recapitulate the complex multi-organ physiology responsible for ABCL homeostasis. While lipoproteins are studied extensively in mammalian models, these systems are not conducive to high-throughput drug discovery. By contrast, the larval zebrafish model system has proven to be a powerful system for *in vivo* drug discovery, as it recapitulates all major aspects of vertebrate physiology in a small, transparent, rapidly developing organism. However, no existing assays are sensitive enough to characterize ABCLs in individual larval zebrafish [15-17], as each larvae contains only a few nanoliters of plasma.

Here we present the LipoGlo reporter as a remarkably sensitive and tractable new tool to study atherogenic lipoproteins. Modern genome engineering techniques were used to fuse the endogenous ApoB gene in zebrafish with an engineered luciferase reporter (NanoLuc), such that each atherogenic lipoprotein would be tagged with a light-emitting molecule. Using this reporter, we were able to develop several independent assays to characterize distinct aspects of the ABCL profile (summarized in Fig. 1a). These include a plate-based assay to measure lipoprotein quantity (LipoGlo-Counting), a gel-based assay to measure lipoprotein size (LipoGlo-Electrophoresis), and chemiluminescent imaging to visualize lipoprotein localization (LipoGlo-Microscopy).

**Figure 1:**
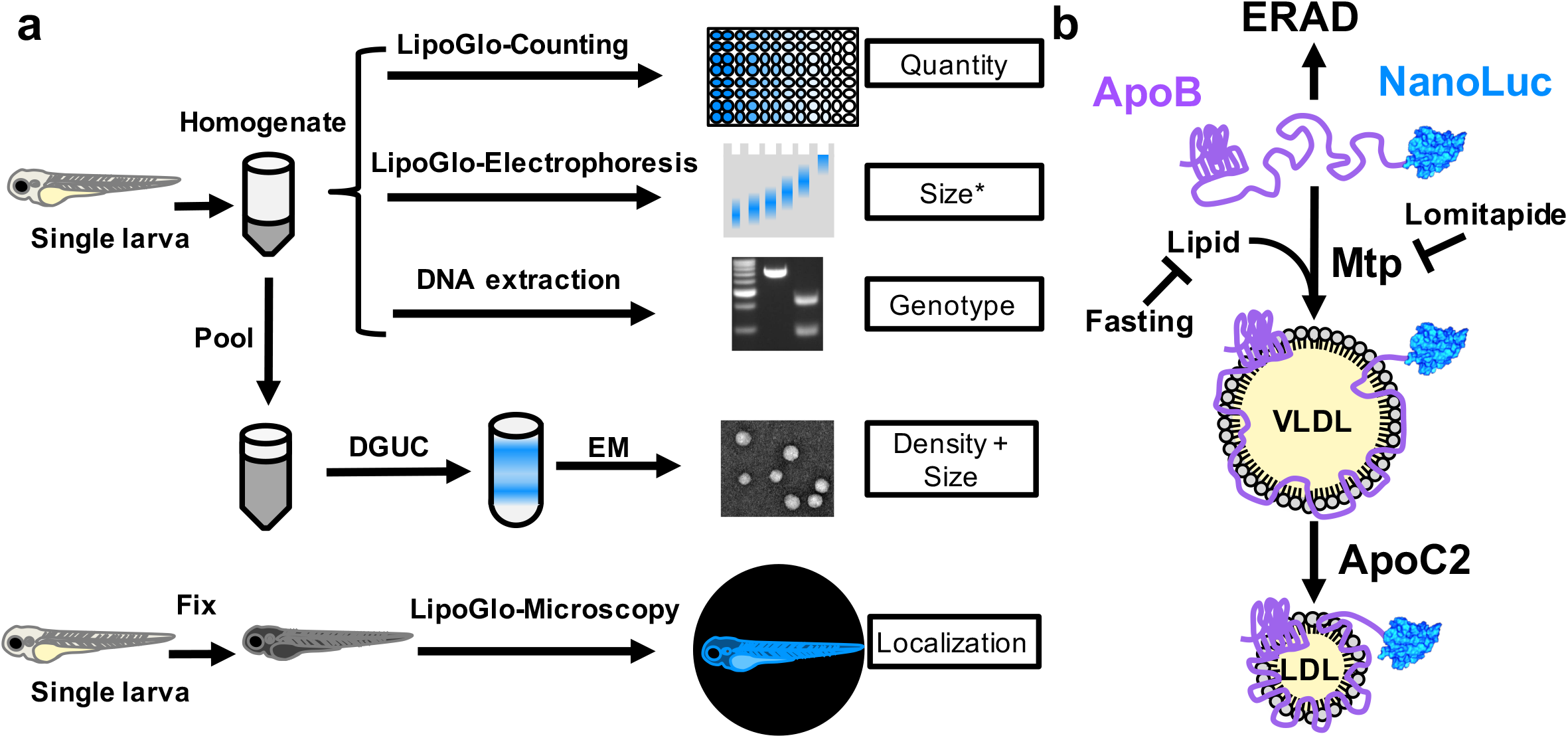
Overview of LipoGlo assays and experimental manipulations. **(a)** Individual larvae carrying the ApoB-NanoLuc reporter are first homogenized in ABCL stabilization buffer. Homogenate can be used for LipoGlo-Counting (a plate-based assay for NanoLuc activity to measure the total number of ABCLs), LipoGlo-Electrophoresis (a Native-PAGE assay to determine the ABCL size/subclass distribution), and DNA extraction for genotyping. Alternatively, lipoprotein density and size can be determined by density-gradient ultracentrifugation on pooled samples (DGUC) followed by electron microscopy. To determine localization of ABCLs *in situ*, individual larvae are fixed in 4% PFA and mounted in low-melt agarose for chemiluminescent imaging (LipoGlo-Microscopy). Asterisk indicates that electrophoretic mobility is an indirect measure of particle size. **(b)** ApoB protein fused to NanoLuc is loaded with lipid through the activity of Mtp to form VLDL particles. In the absence of lipidation, the protein will be rapidly degraded by ERAD. VLDL is lipolyzed by serum lipases that use Apoc2 as an obligate cofactor to produce smaller lipoprotein classes such as LDL. Here we investigate the effects of (i) genetic manipulations (mutations in *mtp* and *apoC2*), (ii) dietary variation (fasting and feeding), and (iii) pharmacological treatment (inhibition of Mtp with lomitapide) on various aspects of the ABCL profile.

We also performed extensive validation of these assays *in vivo* by showing conserved responses to genetic, pharmacological, and dietary manipulations in living zebrafish (summarized in Fig. 1b). Finally, we leveraged the discovery potential of these assays to identify previously uncharacterized associations between ABCLs and the central nervous system [18], as well as identify the poorly characterized gene *pla2g12b* [19] as a potent regulator of lipoprotein particle size that is conserved across vertebrates.

LipoGlo was developed first in larval zebrafish as this organism is uniquely well-suited for high-throughput genetic and small molecule screening, as well as whole-organism imaging. However, LipoGlo represents a highly generalizable tool that can be expanded to function in essentially any organism with atherogenic lipoproteins, and customized with different reporters depending on the research question. This technique has the potential to transform our understanding of atherogenic lipoprotein biology, which may have important clinical repercussions in the treatment of atherosclerotic cardiovascular disease.

## RESULTS

### TALEN-mediated genome engineering enables creation of the LipoGlo reporter

ApoB is an ideal scaffold for creating a reporter of ABCLs. It is both an obligate structural component present in single copy on each lipoprotein particle [20], and is rapidly degraded when not associated with an ABCL via endoplasmic-reticulum-associated protein degradation (ERAD) [21] (Fig. 1b). In mammals there is a single *APOB* gene that can be post-transcriptionally edited into two isoforms: the full-length *APOB-100* expressed primarily in the liver, and the truncated *APOB-48* isoform expressed in the intestine [22, 23]. Although the zebrafish genome contains 3 paralogs of *APOB*, a single paralog (*apoBb.1*) is the clearly dominant isoform, accounting for approximately 95% of the *ApoB* mRNA and protein in larval zebrafish [24]. Known functional elements of ApoB are well conserved in zebrafish, including both the microsomal triglyceride transfer protein (MTP) interacting [25] and LDL-receptor binding [26] domains (Supplementary Fig. 1a). However, the APOB-48 editing site required for production of the truncated (intestine-specific) version of APOB [23] appears to be completely absent in zebrafish (Supplementary Fig. 1b). This creates the opportunity to simultaneously tag both intestine and liver derived ABCLs with a carboxy-terminal fusion to ApoBb.1 in zebrafish.

NanoLuc is an optimized luciferase reporter that generates a quantitative chemiluminescent signal through cleavage of its substrate molecule, furimazine [27]. This reporter is remarkably bright (~100 times brighter than firefly luciferase), small (19.1 kDa), stable, and provides robust signal to noise ratios that enable accurate detection even at femtomolar concentrations [27]. The NanoLuc coding sequence was introduced as a carboxy-terminal fusion to the endogenous ApoBb.1 gene in zebrafish through homology directed repair of a double-stranded break [28]. Capped mRNA encoding a TALEN pair targeting the ApoBb.1 stop codon was co-injected with a donor DNA construct to induce homology-directed repair. The donor construct contains the NanoLuc coding sequence flanked on either side by several hundred base pairs of sequence homologous to the genomic sequence upstream and downstream of the ApoBb.1 stop codon (Supplementary Fig. 2). Injected embryos were raised to adulthood and their progeny were screened for NanoLuc activity and subsequently for error-free integration at the target locus. The resulting tagged lipoproteins are quantified using the Nano-Glo assay (Promega Corp., N1110), which led us to name this system LipoGlo.

Fish homozygous for the LipoGlo reporter are healthy, fertile, and do not display any abnormal morphological or behavioral phenotypes. Additionally, larvae homozygous for the LipoGlo reporter show a two-fold increase in LipoGlo signal relative to their heterozygous siblings (Supplementary Fig. 2c). Together, these data suggest that the LipoGlo reporter does not disrupt normal production, secretion, and turnover of lipoprotein particles.

### LipoGlo-Counting reveals changes in ABCL abundance

The LipoGlo-Counting method uses a 96-well plate based assay to detect NanoLuc activity and quantify ABCL abundance. In order to validate the LipoGlo reporter and evaluate the degree of similarity between zebrafish and mammalian lipoprotein homeostasis, the lipoprotein profile was assayed across development, as well as in response to genetic, pharmacological, and dietary manipulation (Fig. 1b). Individual larvae carrying the LipoGlo reporter are homogenized in a standard volume of ABCL stabilization buffer (100 μL) using either a pellet pestle for low throughput sample processing (Fisher scientific, 12-141-363), or a microplate horn sonicator for processing of 96 samples simultaneously in plate format (QSonica, 431MPX). The ABCL stabilization buffer contains protease inhibitors, pH buffers, and cryoprotectant to ensure sample stability during processing and storage. A portion of the homogenate (40 μL) is mixed with an equal volume of Nano-Glo assay buffer and quantified in a plate reader. The remaining homogenate is either stored frozen for later use, or used for additional assays (Fig. 1a).

ABCL levels were measured throughout development from 1 – 6 days post-fertilization (dpf) using zebrafish carrying the LipoGlo reporter in the wild-type (WT) genetic background (Fig. 2a). During this window of development, embryos are in the lecithotropic (yolk-metabolizing) stage [29]. All nutrients required for development are provided by the maternally deposited yolk, until the yolk becomes depleted between 5 and 6 dpf and the larvae begin to rely on exogenous food. Yolk lipid is packaged into ABCLs by the yolk syncytial layer (YSL), a specialized embryonic organ that expresses many genes involved in ABCL production including *ApoBb.1* [24]. Accordingly, ABCL levels are quite low early in development, but increase between 1 – 3 dpf as more yolk lipid is packaged into ABCLs (Fig. 2a). As the larvae are not provided with food, ABCL levels drop later in development as rates of lipoprotein metabolism and turnover exceed rates of production following yolk depletion.

**Figure 2:**
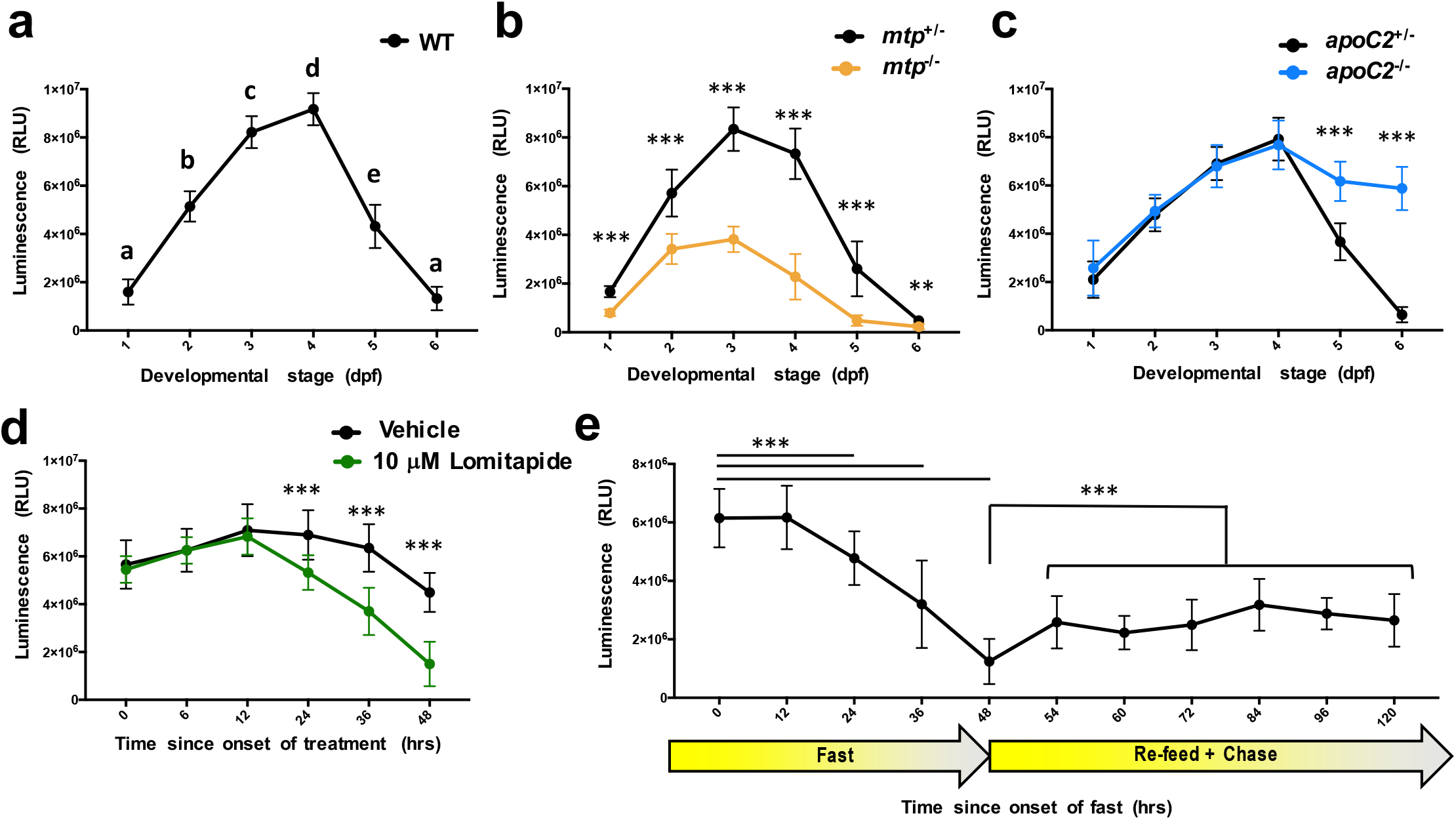
LipoGlo-Counting reveals conserved ABCL responses to genetic, dietary, and pharmacological stimuli. **(a)** LipoGlo signal throughout WT larval zebrafish development (1 – 6 dpf). Time points designated with different letters are statistically significantly different (n=24, ANOVA p<0.0001,Tukey’s HSD p<.0001). **(b)** Comparison of LipoGlo signal between *mtp^-/-^* mutants (defective in lipoprotein synthesis) and *mtp^+/-^* siblings during larval development (n≈16, Two-way robust ANOVA p<0.0001 for genotype and stage, Games-Howell p<.001). **(c)** Comparison of LipoGlo signal between *apoC2^-/-^* mutants (defective in lipoprotein breakdown) and *apoC2^+/-^* siblings during larval development (n≈12, Two-way robust ANOVA p<0.0001 for genotype and stage, Games-Howell p<.0001). **(d)** Effect of lomitapide (10 μM, Mtp inhibitor) on LipoGlo signal (3 – 5 dpf) (n=30, Two-way robust ANOVA p<0.000lfortreatment and time, Games-Howell p<.0001). **(e)** LipoGlo levels were measured overtime throughout a fast, re-feed, and chase period. Larvae were fed a standard diet *ad libitum* from 5 to 10 dpf, and then were deprived of food for 48 hours (fast period). Larvae were then fed a high-fat (5%egg yolk) diet for 6 hours, transferred to fresh media, and assayed at various time points for the 72 hours following the onset of feeding (48-120 hours) (n=30, Welch’s ANOVA p<0.0001, Games-Howell p<.0001). Results represent pooled data from three independent experiments, “*n*” denotes number of samples per data point.

LipoGlo reporter fish were then crossed with fish harboring mutations in essential components of the ABCL production and breakdown pathways. Microsomal Triglyceride Transfer Protein (Mtp) is responsible for loading nascent ApoB with lipid to form ABCLs [30], and Apolipoprotein-C2 (ApoC2) is a cofactor for lipoprotein lipolysis [31] (outlined in Fig. 1b). As expected, *mtp^-/-^* mutants [32] exhibit profound defects in ABCL production detectable from the earliest stages of development (Fig. 2b). By contrast, *apoC2^/-^* mutants [15] produce lipoproteins normally but show significantly reduced levels of particle breakdown and turnover compared to sibling controls (Fig. 2c).

To probe the effects of transient Mtp inhibition on larval lipoprotein homeostasis, larvae were exposed to lomitapide. Lomitapide is a pharmaceutical inhibitor of Mtp used to treat familial hypercholesterolemia in humans [33]. Larvae were treated with 10 μM lomitapide or vehicle control for 48 h (3-5 dpf), and treated larvae showed a more rapid decline in NanoLuc levels than vehicle-treated controls. This observation is consistent with lomitapide inhibiting ABCL production and leading to an accelerated decline of ApoB-NanoLuc levels (Fig. 2d).

To test the effect of food intake on ABCL levels, larvae were subjected to a fasting and re-feeding experimental paradigm. Larvae were fed a standard diet (Gemma 75, Skretting USA) for 5 days (from 5-10 dpf) to adapt to food intake and reach a physiologically relevant baseline level of ABCLs. Following the initial feeding period, larvae were fasted for 48 h (sampled every 12 h), re-fed with a high-fat meal of 5% egg-yolk [34], and sampled at various time points after the meal (the chase period). ApoB-NanoLuc levels were stable for the first 12 h of the fast, but declined rapidly for the duration of the fasting period (Fig. 2e, 0-48 h). Following the high fat meal (6 h of feeding from time point 48 h to time point 54 h), there was an immediate increase in ApoB-NanoLuc levels (Fig. 2e, 48-120 hrs). ApoB-NanoLuc levels did not recover to their pre-fasted state following the high-fat meal, but rather remained at an intermediate level for a prolonged period (the duration of the chase period, 72 h).

### Determination of lipoprotein size distribution using LipoGlo-Electrophoresis

There are numerous classes of ABCLs, many of which can be differentiated based on particle size [35]. Native polyacrylamide gel electrophoresis (Native-PAGE) has previously been used to separate ABCLs based on size, but requires a relatively large volume of plasma (25 μL) stained with lipophilic dyes [36]. The LipoGlo-Electrophoresis method subjects crude larval homogenate (containing only nanoliters of plasma) to Native-PAGE to separate lipoproteins, followed by in-gel detection of NanoLuc activity. To analyze the ABCL size distribution over development and in response to genetic, pharmacological, and dietary manipulations, representative frozen aliquots of larval homogenate from a given condition were thawed on ice. A portion of the thawed homogenate (12 μL) was mixed with 5x loading dye (3 μL) and separated via Native-PAGE (3% gel for 275 Volt-h). Following separation, the glass front plate was removed to expose the gel surface, and 1 mL of TBE containing Nano-Glo substrate solution (2 μL) was added to the plate and spread evenly using a thin plastic film. The gel was then imaged using the Odyssey Fc chemiluminescent detection system (LI-COR Biosciences). Together, this protocol is referred to as LipoGlo electrophoresis.

Smaller lipoproteins are expected to migrate further into the gel, and larger lipoproteins to show concomitantly less mobility (Fig. 3a). Following electrophoretic separation, ABCLs can be divided into four different classes based on their migration distance. ABCLs that remain within the loading well are classified as the “zero mobility” (ZM) fraction, which should include chylomicrons [37], remnants, aggregates [38], and intracellular ApoB complexed with components of the secretory pathway (such as the ER, golgi, and other secretory vesicles) [39]. Species that do migrate into the gel are classified as either Very Low-Density Lipoproteins (VLDL), Intermediate-Density Lipoproteins (IDL), or Low-Density Lipoproteins (LDL) based on their electrophoretic mobility. Di-I-Labeled fluorescent LDL (L3482, ThermoFisher Scientific) is used as a migration standard to ensure consistent classification of ABCL species between gels, with the migration distance of this species corresponding to one ladder unit (L.U.). Although the Di-I stain significantly reduces electrophoretic mobility of the human LDL and therefore does not align with NanoLuc-labeled LDL (data not shown), this band provides a highly reproducible standard for registration and normalization across gels (Supplementary Fig. 3).

**Figure 3:**
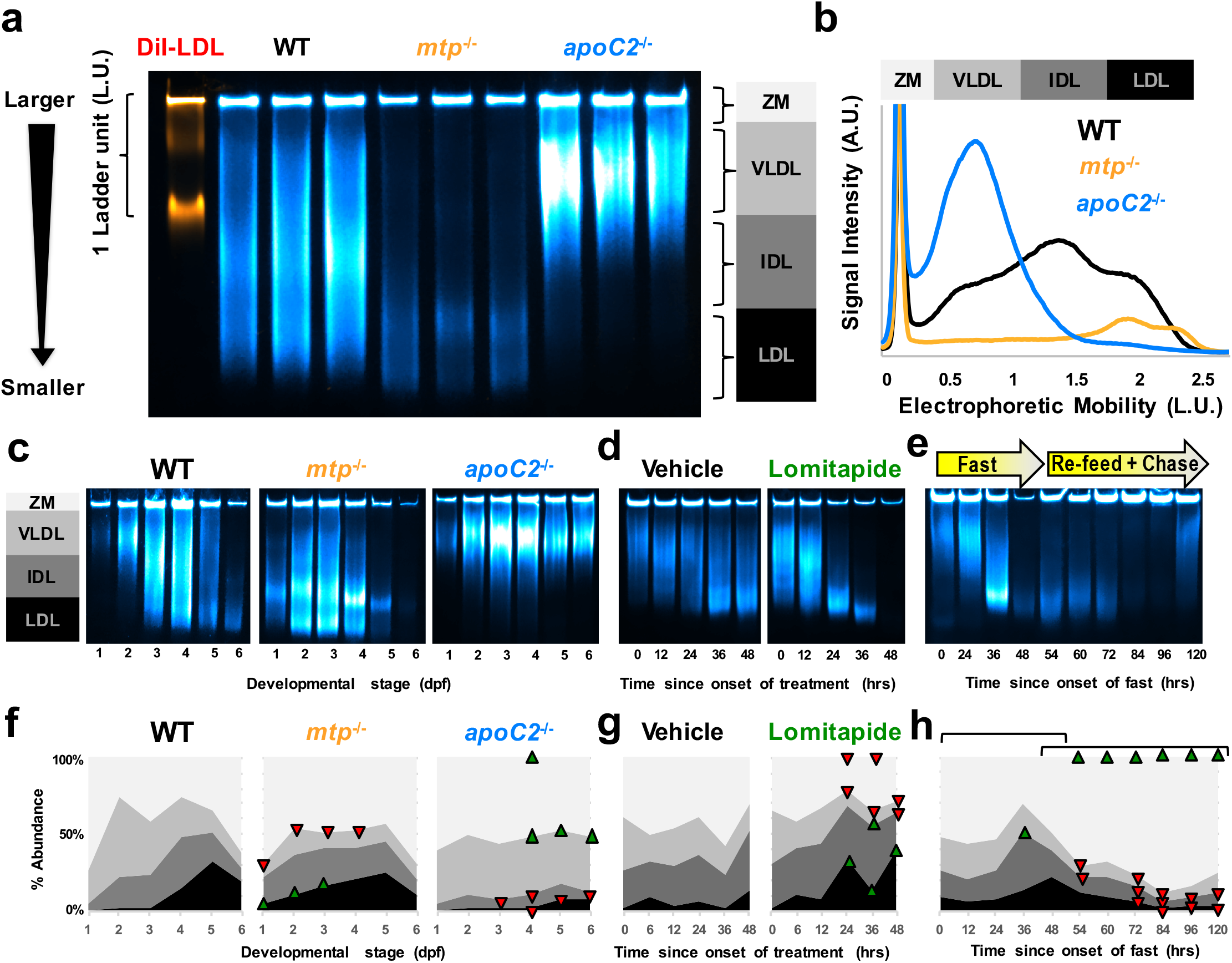
Changes in lipoprotein size distribution revealed through LipoGlo-Electrophoresis. **(a)** Representative image of the fluorescent Dil-LDL migration standard and LipoGlo emission from WT, *mtp^-/-^*, and *opoC2^-/-^* genotypes (4 dpf). ABCLs are divided into 4 classes based on their mobility, including ZM (zero mobility) and three classes of serum ABCLs (VLDL, IDL, LDL). Image is a composite of chemiluminescent (LipoGlo, blue) and fluorescent (Dil-LDL, orange) exposures. Gel is a representative image from one of the three independent experiments performed. **(b)** Vertical plot profile generated in ImageJ from gel image displayed in (a), note that the ZM peak has been appended to highlight differences in serum lipoprotein classes. **(c-e)** Representative gel images (one of three independent experiments shown) and **(f-h)** pooled LipoGlo-electrophoresis quantification data from larval lysates used in Figure 2. Relative abundance of subclasses is color-coded as shown in (a). Upward-facing arrowheads (green) indicate significant enrichment of that species at that time point compared to controls, and downward-facing arrowheads (red) indicate depletion **(f)** Subclass abundance at each day of larval development in WT (n=9, Welch’s ANOVA p<0.0001 for each subclass over time), *mtp^-/-^* (n=9, Two-way robust ANOVA p<0.001 for VLDL and LDL, Games-Howell p<.01), and *opoC2^-/-^* (n=9, Two-way robust ANOVA p<0.01 for all classes, Games-Howell p<.005) genetic backgrounds. **(g)** Subclass abundance from 3-5 dpf in larvae treated with 10μM lomitapide or vehicle control (n=9, Two-way robust ANOVA p<0.001 for all classes except IDL, Games-Howell p<.01). **(h)** Subclass abundance from 10-15 dpf in larvae subjected to a fasting and re-feeding paradigm. The first bracket delineates changes relative to time 0 (the onset of the fasting period), and the second bracket delineates changes relative to time point 48 (the onset of the re-feeding period) (n=9, Welch’s ANOVA p<0.0001 for each subclass over time, Games-Howell p<.01). Supplementary Figure 4 displays standard deviations for panels f-h. Results represent pooled data from three independent experiments, “*n*” denotes number of samples per data point.

In order to define physiologically relevant migration boundaries between ABCL classes, ABCL profiles were compared for WT, *mtp^-/-^*, and *apoC2^-/-^* mutant lines at 4 dpf (Fig. 3a,b). *ApoC2^-/-^* mutants are unable to lipolyze VLDL, which allowed us to define the VLDL bin from .3 – 1 ladder units. Conversely, *mtp*^-/-^ mutants display a bimodal peak of small LDL-like particles at this stage of development, which was used to define the LDL bin as 1.7 – 2.4 ladder units from the origin. Wild-type larvae have a peak of intermediate-sized lipoproteins at this stage, which corresponds to the IDL region from 1 – 1.7 ladder units. ABCLs migrating less than .3 ladder units were considered to be in the zero-mobility fraction (ZM) (Fig. 3a).

Gel images were transformed into plot profiles in ImageJ for quantification (Fig. 3b). The provided Gel Quantification Template (supplementary file 1) contains instructions and formulas for automatically calculating bin cutoffs for each ABCL class based on the migration of the Di-I standard and quantifying the relative intensity of each bin. To visualize the distribution of ABCL classes over time, each species was color coded with darker colors corresponding to smaller lipoproteins and plotted as an 100% stacked area chart (Fig. 3f-h). Upward-facing green arrowheads or downward-facing red arrowheads are used to indicate which species show significant enrichment or depletion (respectively) relative to the control group (Fig. 3f-h). Additional plots were generated that present these data grouped by ABCL class (rather than genotype) (Supplementary Fig. 4).

Using LipoGlo-Electrophoresis over the course of zebrafish larval development revealed that in the early embryonic stages (1-2 dpf), the wild-type ABCL profile is dominated by VLDL (Fig. 3c,f), which are directly produced by the YSL. By 3 and 4 dpf, these VLDL particles have been lipolyzed to generate the smaller IDL and LDL classes. When the maternal yolk has been depleted (5-6 dpf) and in the absence of exogenous food, VLDL production is significantly attenuated as indicated by the enrichment of the small lipolyzed lipoproteins. The ABCL profile dynamics are much more static in the *mtp^-/-^* and *apoC2^-/-^* mutant lines. *mtp^-/-^* mutants produce smaller IDL and LDL-like particles from the earliest stages of development, and *apoC2^-/-^* mutants show a VLDL peak that persists throughout development (Fig. 3c,f). Consistent with the data from *mtp^-/-^* mutants, pharmacological treatment with a potent MTP inhibitor (Lomitapide) effectively blocks the production of new VLDL particles (Fig. 3d,g) leading to the accumulation of lipolyzed species such as IDL and LDL.

Consistent with the mammalian literature, a robust post prandial response to a high lipid meal (egg yolk emulsion) was observed in the distribution of ABCL subclasses of larval zebrafish (Fig. 3e,h). After fasting (48 h), there is significant depletion of VLDL and enrichment of LDL, consistent with cessation of VLDL production due to limited nutrient availability. A subsequent high-fat meal produces a significant increase in the ZM band, and progressive depletion of LDL.

### Electrophoretic mobility correlates well with lipoprotein density and size

Electrophoretic mobility in Native-PAGE is a function of both size and charge, so it is important to evaluate whether differences in migration truly reflect different lipoprotein sizes or if they are the result of differentially charged lipoproteins. Density gradient ultracentrifugation (DGUC) is the gold standard for discerning different subclasses of ABCLs, as larger lipoprotein classes are more buoyant resulting from their large lipid core. To evaluate concordance between DGUC and the LipoGlo assays, we developed a DGUC protocol (based on the method described by Yee et al., [40]) to separate pooled larval homogenate into density fractions. We then subjected fractions to (i) LipoGlo electrophoresis to characterize their electrophoretic mobility, (ii) a plate read assay to quantify ApoB-NanoLuc levels, and (iii) negative-staining electron microscopy to visualize particle size directly [41] (Fig. 4). Importantly, denser fractions showed higher electrophoretic mobility and smaller particle sizes across all genotypes, demonstrating that electrophoretic mobility is a reliable method for differentiating ABCL classes and can be used as a proxy to estimate particle size and density.

**Figure 4:**
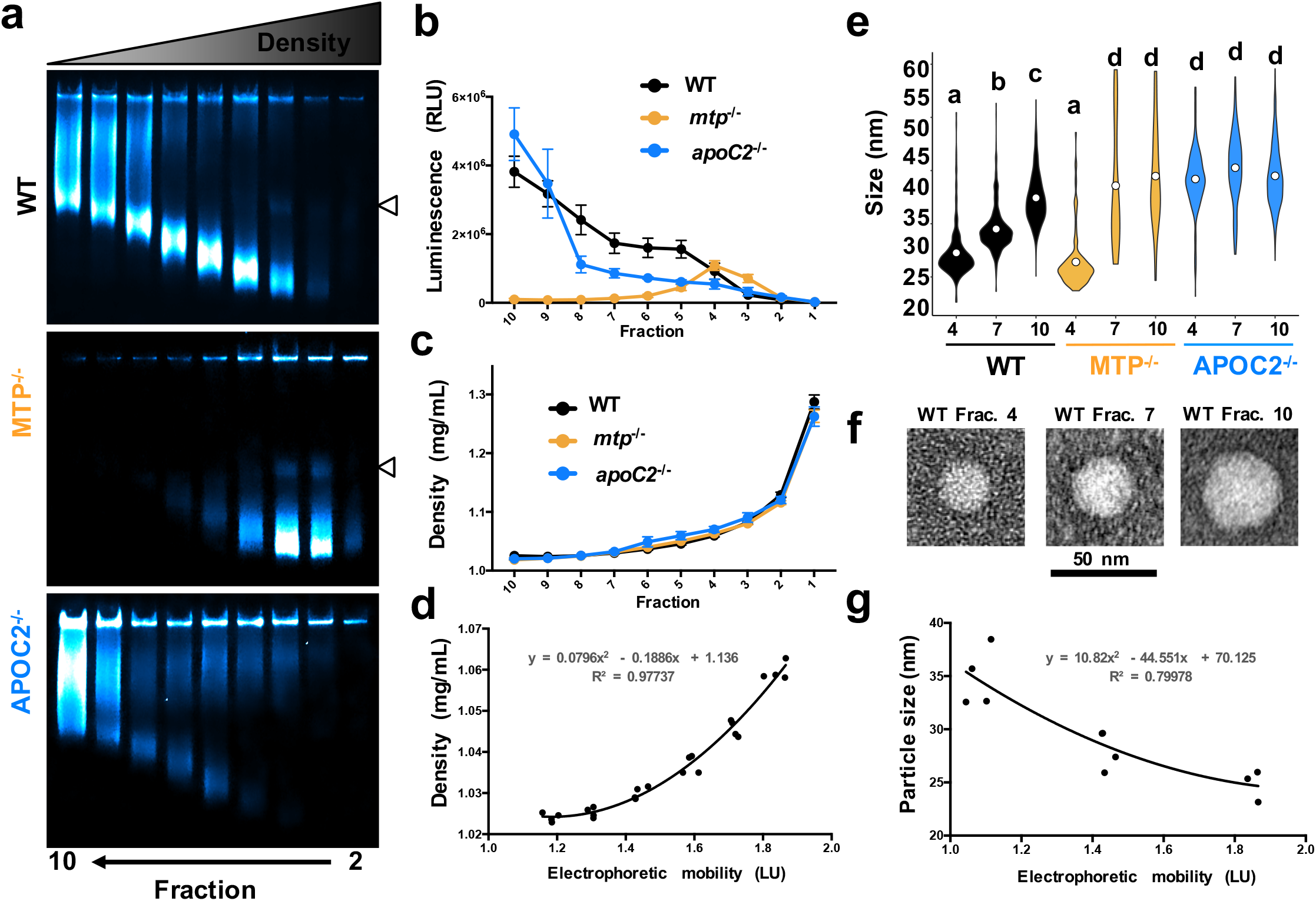
Concordance between LipoGlo electrophoresis and classical ABCL size characterization techniques. DGUC was performed on pooled larval homogenate (4 dpf) from WT, *mtp^-/-^*, and *apoC2^-/-^*, and separated into 10 equal fractions of approximately 500 μL each by drip-elution (dense bottom fractions eluted first). **(a)** Fractions 2-10 were subjected to Native-PAGE, and denser fractions showed higher electrophoretic mobility. Some fractions show a faint lower mobility band (indicated at right by white arrowhead), possibly indicative of lipoprotein dimerization. **(b)** A plate-based assays of NanoLuc activity revealed the expected enrichment of VLDL in *apoC2^-/-^* mutants, and enrichment of LDL in *mtp^-/-^* mutants (confirming results reported in Fig. 3b). **(c)** A refractometer (Bausch and Lomb) was used to determine the refractive index of each fraction and density was calculated via the formula D = 3.3508 x RI - 3.4675. DGUC showed highly reproducible density profiles between replicates and genotypes. **(d)** The density of WT fractions 4 - 9 was plotted as a function of peak electrophoretic mobility for that fraction, and the second order polynomial function (y = 0.0796×2 - 0.1886x + 1.136) was able to represent this relationship with remarkable accuracy (R^2^ = 0.97737) indicating that electrophoretic mobility is a useful proxy for lipoprotein density. **(e)** Fractions 4,7, and 10 were subjected to negative-staining electron microscopy to directly visualize the size of particles in each fraction. I n the wild type samples, the average particle diameterwas 24.7±5.6,29.0±4.1, and 34.9±4.7 nm for fractions 4,7, and 10 respectively. There was no significant difference in particle size between fraction 4 of the wild-type and *mtp^-/-^* mutant samples (average diameter of 23.2 ±6.6 nm). Particles were nearly undetectable in fractions 7 and 10 in the *mtp^-/-^* mutant sample so particle diameter shows enormous variability. ABCLs in each *apocC2^-/-^* mutant fraction were significantly larger than all WT fractions, with diameters of 39.0±8.0, 40.9±7,2, and 39.1±5.9 nm respectively (n≈170, Welch’s ANOVAp<0.0001, Games-Howell p<.0001). **(f)** Representative images of lipoproteins from the three wild-type fractions are shown. **(g)** The second-order polynomial function y= 10.82×2 - 44.551x + 70.125 approximated the relationship between electrophoretic mobility and density in wild-type samples with reasonable accuracy (R^2^ = 0.79978). Results represent pooled data from four independent experiments.

### LipoGlo-Microscopy reveals whole-organism ABCL localization

The transparency of larval zebrafish offers the unique opportunity to perform whole-mount imaging, which has enabled us to perform the first characterization of changes in ABCL localization throughout an intact organism. The same developmental, genetic, dietary, and pharmacological manipulations described above (Figs. 2-3) were performed, but rather than being homogenized, larvae were fixed in 4% paraformaldehyde (PFA) for 3 h at room temperature, washed in PBS-tween, and mounted in low-melt agarose [42] supplemented with Nano-Glo substrate solution. Mounted larvae were imaged in a dark room on a Zeiss Axiozoom V16 equipped with a Zeiss AxioCam MRm set to collect a single brightfield exposure followed by multiple exposures with no illumination (chemiluminescent imaging).

The differences between WT, *mtp^-/-^*, and *apoC2^-/-^* mutants were most apparent at 6 dpf (Fig. 5a). At this stage, the yolk is depleted and larvae are in a fasted state as no exogenous food has been provided. In WT larvae, signal is quite low throughout the body, but is clearly visible in the lipoprotein-producing tissues (liver and intestine). We observed a previously undescribed association of ApoB with the spinal cord (Fig. 5b, and Supplementary Fig. 5a) as evidenced by colocalization with the central-nervous system marker *Tg(Xla.Tubb2:mApple-CAAX*). This reporter uses the tubulin beta-2 promoter from *X. Laevis* to drive a membrane-targeted mApple fluorophore specifically in the CNS. A dorsal view revealed enrichment of NanoLuc signal in particular brain regions (Fig. 5c), which we hypothesize may correspond to the brain ventricle. In *mtp^-/-^* mutants, ApoB is very low outside of the lipoprotein-producing tissues, consistent with defects in loading ApoB with lipid to form a secretion-competent ABCL. *ApoC2^-/-^* mutants show remarkably high signal throughout the body, consistent with their inability to process and turnover lipoproteins (Fig. 5a).

**Figure 5:**
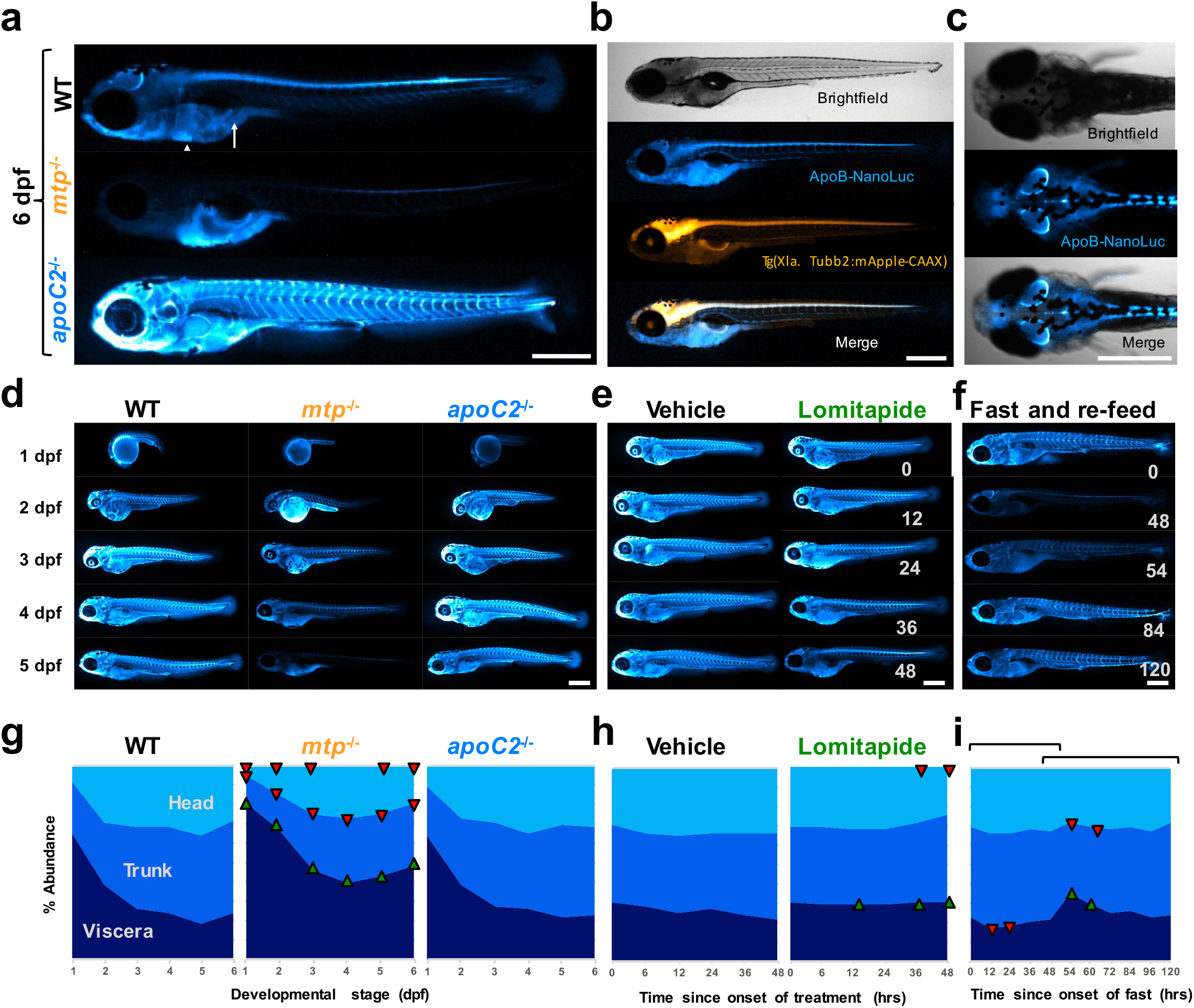
Whole-mount imaging of ABCL localization using LipoGlo chemiluminescent microscopy. **(a)** Representative images of ABCL localization patterns from analysis of 15 larvae per genotype from WT, *mtp^-/-^*, and *apoC2^-/-^* genotypes (6 dpf). The white arrow and arrowhead mark the larval intestine and liver respectively. **(b)** LipoGlo signal colocalizes with the central nervous system marker *Tg(Xla. Tubb2-mApple-CAAX*), quantification in Supplementary Figure 5. **(c)** LipoGlo signal localized to subregions of the CNS. **(d-f)** Representative images and **(g-i)** quantification of ABCL localization across developmental, genetic, pharmacological, and dietary manipulations. Upward-facing arrowheads (green) indicate significant enrichment of that species at that time point compared to controls, and downward-facing arrowheads (red) indicate depletion. **(g)** Signal localization at each day of larval development in WT (n=15, Welch’s ANOVA p<0.0001 for each region over time), *mtp^-/-^* (n=15, Two-way robust ANOVA p<0.001 for all regions, Games-Howell p<0.001), and *apoC2^-/-^* (n=15, Two-way robust ANOVA was not significant for any region) genetic backgrounds. **(h)** Signal localization from 3-5 dpf in larvae treated with 10μM lomitapide or vehicle control (n=15, Two-way robust ANOVA p<0.001 for head and viscera, Games-Howell p<.0001). **(i)** Subclass abundance from 10-15 dpf in larvae subjected to a fasting and re-feeding paradigm. The first bracket delineates changes relative to time 0 (the onset of the fasting period), and the second bracket delineates changes relative to time point 48 (the onset of the re-feeding period) (n=15, Welch’s ANOVA p<0.0001 for each region, Games-Howell p<.005). Supplementary Figure 4 displays standard deviations for panels g-i. Results represent pooled data from three independent experiments, “n” denotes number of samples per data point. Body regions were defined as outlined in Supplementary Figure 5. Scale bars = 500 μm.

Images were quantified by creating separate regions of interest for the viscera, trunk, and head regions (Supplementary figure 5c) and comparing the relative levels of NanoLuc signal in each of these areas. During development, signal was initially highly enriched in the visceral region, which contains the yolk and YSL, and then gradually increases in the trunk and head regions (Fig. 5d,g). This is consistent with the vectorial transport of lipid from the YSL to the circulatory system and peripheral tissues. The distribution of ApoB between these three regions was not significantly changed in *apoC2^-/-^* mutants, whereas *mtp*^-/-^ mutants showed enrichment in the viscera and depletion in the peripheral tissues at all time points (Fig. 5d,g). Results were also grouped by region to facilitate comparison of each class between genotypes (Supplementary Fig. 4).

### LipoGlo assays reveal Pla2g12b as an important regulator of ABCL homeostasis

In an effort to identify novel regulators of the ABCL profile using the LipoGlo system, we screened through several mutant lines from the zebrafish mutation project [43] that had predicted mutations in genes involved in lipid metabolic pathways. Larvae homozygous for an essential splice site mutation (sa659) in phospholipase A2 Group XII B (*pla2g12b*) showed perturbations in their ABCL profile (Fig. 6). Homozygous mutant larvae exhibited lower levels of ApoB at multiple stages (Fig. 6a), and also appeared to have defects in lipoprotein secretion as evidenced by enrichment of visceral ApoB-NanoLuc levels (Fig. 6b,e). However, the most striking defect in *pla2g12b^-/-^* mutant larvae is a pronounced change in the ABCL size distribution. Even at 1 dpf, significant accumulation of small lipoproteins in the size range of LDL, and depletion of the larger particle classes, were evident (Fig. 5c,d).

**Figure 6:**
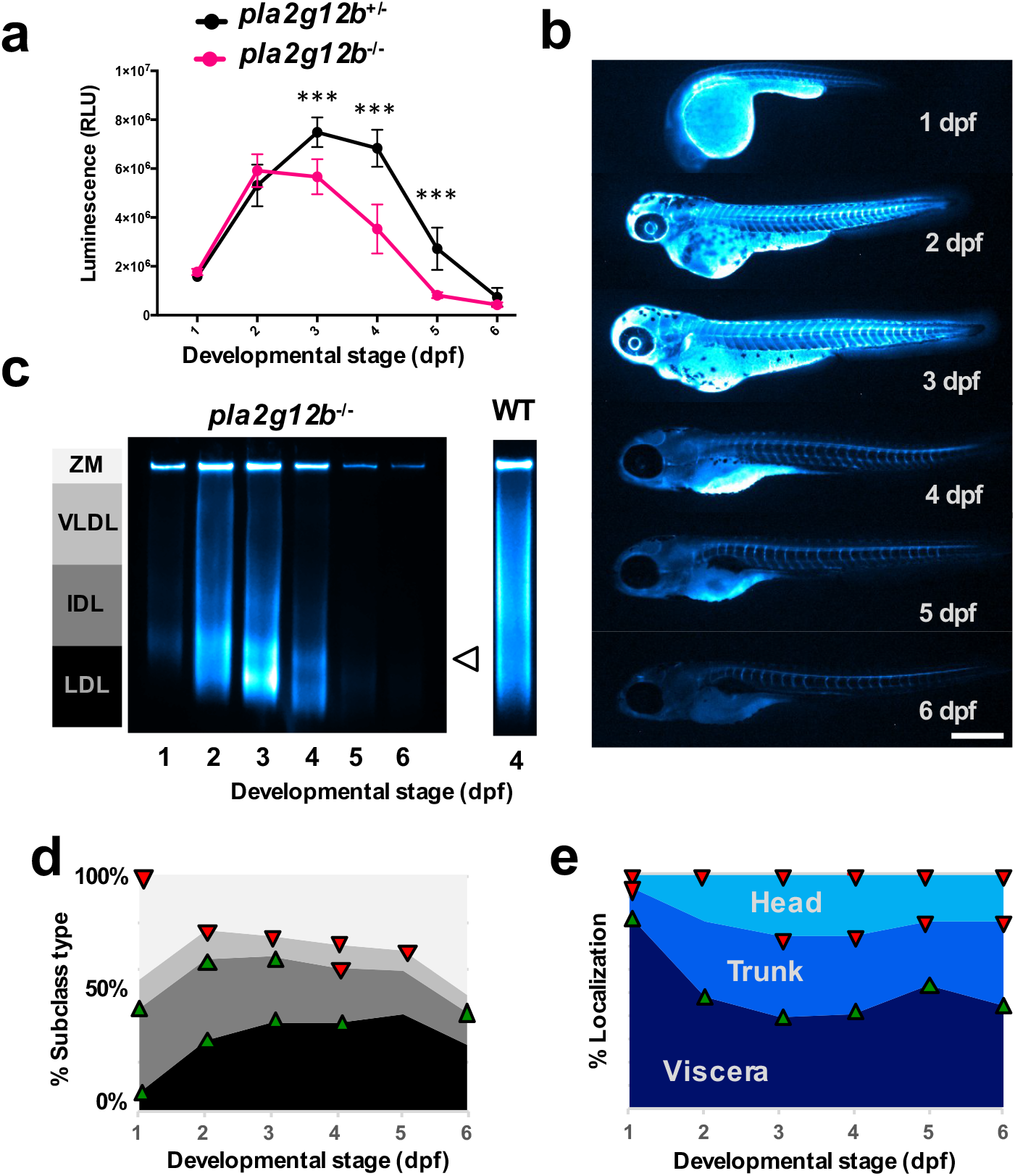
LipoGlo reveals profound alterations in the ABCL profile in *pla2g12b^-/-^* mutant larvae. **(a)** Comparison of LipoGlo signal between *plo2gl2b^-/-^* and *plo2gl2b^+/-^* siblings during larval development (1 – 6 dpf) (n≈ll, Two-way robust ANOVA p<0.0001 for genotype and stage, Games-Howell p<.0001). **(b)** Representative images (n=15) of ABCL localization collected by LipoGlo chemiluminescent imaging throughout development (1 – 6 dpf), and **(e)** quantification of percent localization into previously described subregions (n=15, Two-way robust ANOVA p<0.001 for all regions, Games-Howell p<0.0001). **(c)** Representative gel (n=4) showing production of abnormally small lipoproteins (white arrowhead) with a lane of the 4 dpf WT profile (reproduced from Fig. 3a) provided for reference. **(d)** Quantification of LipoGlo emission pattern from LipoGlo electrophoresis in *pla2gl2b^-/-^* larvae (1 – 6 dpf). Upward-facing arrowheads (green) indicate significant enrichment of that species at that time point compared to WT, and downward-facing arrowheads (red) indicate depletion (n=9, Two-way robust ANOVA p<0.001 for all species, Games-Howell p<.01). Results represent pooled data from three independent experiments, “n” denotes number of samples per data point. Body regions were defined as outlined in Supplementary Figure 5. Scale bars = 500 μm.

## DISCUSSION

### The LipoGlo reporter does not disrupt lipoprotein homeostasis

When generating a fusion protein, it is essential to evaluate whether introduction of the tag disrupts native protein function. This is particularly important in the case of tagged lipoproteins, as these particles have a complex life cycle that involves interaction with numerous cell and tissue types. To evaluate whether the NanoLuc tag disrupted lipoprotein homeostasis, LipoGlo larvae were subjected to various genetic, dietary, and pharmacological manipulations known to affect the lipoprotein profile. Detailed characterization of the lipoprotein profile in response to each of these stimuli revealed the expected results in each case. These results validate that NanoLuc-tagged lipoproteins exhibit all of the central hallmarks of endogenous ABCLs, including MTP-dependent maturation, APOC2-dependant lipolysis, responsiveness to nutrient availability, and expected density and size distributions.

These observations establish a precedent for the use of ApoB-fusion proteins as a sensitive and specific approach to monitor atherogenic lipoproteins *in vivo*. This approach can now be generalized to function in essentially any model system expressing ApoB, and expanded to use alternative tags such as fluorescent reporters for high-resolution imaging or affinity tags to study the lipoprotein interactome. Potential applications of LipoGlo thus extend well beyond the study of lipoprotein abundance, size, and localization using zebrafish.

### LipoGlo assays show excellent concordance with existing methods

Atherogenic lipoproteins are the primary drivers of atherosclerotic cardiovascular disease, which has made lipoprotein profiling an essential technique in both clinical and research settings. Existing methods to characterize ABCLs generally rely on large amounts of starting material (microliters of plasma), use indirect measurements (e.g. lipid profiling), require expensive equipment and specialized training (e.g. an ultracentrifuge or electron microscope), have a relatively low throughput capacity, and are restricted to studying plasma lipoproteins. The LipoGlo reporter circumvents all of these issues and engenders unprecedented speed, sensitivity, and tractability to the study of the lipoprotein profile. Importantly, these assay shows excellent concordance with traditional methods, as evidenced by the tight correlation between particle size estimates measured by LipoGlo electrophoresis and both density gradient ultracentrifugation and negative-staining electron microscopy.

One limitation of the LipoGlo electrophoresis method reported here is that it is unable to resolve ABCLs above a certain size threshold, which are clustered together as the zero-mobility fraction. Further research will thus be required to conclusively determine which lipoprotein species are present within this band. For example, this band is highly enriched in response to a high-fat meal. This observation is consistent with enrichment of intracellular ApoB and the largest lipoprotein classes (chylomicrons and remnants), both of which should increase in response to high-fat diet. It is interesting to note, however, that when lipoproteins are processed through tandem density gradient ultracentrifugation and LipoGlo-electrophoresis, there is significant signal in the ZM fraction even in the higher density fractions. This indicates that the ZM band is not exclusively composed of very large triglyceride-rich lipoproteins, but also includes additional denser species. We suspect this may reflect the physiological level of lipoprotein aggregation, which has been implicated as yet another determinant of cardiovascular disease risk.

### The larval zebrafish is a powerful new system to study lipoprotein biology

The LipoGlo system is significantly more sensitive than existing lipoprotein characterization assays. To illustrate this, we have performed extensive lipoprotein characterization on individual zebrafish larvae. A single larva contains approximately one-thousand times less plasma (nanoliters rather than microliters) than is traditionally used for lipoprotein profiling, yet this is more than enough material to run multiple LipoGlo assays. The larval zebrafish has been used extensively to study metabolic diseases including cardiovascular disease [44], but the inability to characterize the atherogenic lipoprotein profile in this model presented a significant limitation. LipoGlo enabled comprehensive characterization of the atherogenic lipoprotein profile throughout larval development, which will serve as a valuable resource for the zebrafish research community. LipoGlo was also used to demonstrate numerous similarities in lipoprotein processing between larval zebrafish and humans.

The larval zebrafish is also an unparalleled vertebrate system for high-throughput screening. The simple plate-based LipoGlo-counting method enables processing of tens of thousands samples a day, and is thus readily conducive to high-throughput genetic and small-molecule screening. The LipoGlo-electrophoresis and imaging protocols can also achieve respectable throughput capacity on the order of hundreds of samples per day, and can thus be used as tractable secondary screening assays or for stand-alone small-scale screens. The confluence of LipoGlo and the zebrafish model system create the first opportunity to perform unbiased screens for novel modulators of ABCLs, enabling powerful unbiased discovery approaches to be applied to the field of lipoprotein biology.

### Sensitive lipoprotein profiling may provide new insights into abetalipoproteinemia

Human mutations in the *mtp* gene result in severe reduction or complete lack of ABCLs, a disease called abetalipoproteinemia. The *mtp^stl^* allele studied contains a single missense mutation in a highly conserved residue (L475P) [32]. Although this is thought to result in production of a nonfunctional protein, a true null allele would be expected to result in a complete lack of ABCLs. The *mtp^stl^* homozygous mutants are unequivocally able to produce and secrete a ABCLs (although they are smaller and less abundant) early in development, although at the later stages ABCLs are essentially undetectable. These observations suggest that the *mtp^stl^* allele is either a strong hypomorph, or that ABCLs can be produced without the activity of Mtp. The complete lack of ABCLs later in development occurs once rates of particle turnover and uptake begin to greatly exceed rates of production. This observation highlights the LipoGlo system as a useful tool to study allelic series of *mtp*. By studying different alleles at the earliest stages of development when rates of lipoprotein turnover are low, it may be possible to distinguish between true null alleles and varying degrees of hypomorphic mutations. Such information would be useful in predicting the severity of different abetalipoprotienemia mutations in humans.

Additionally, the *mtp^-/-^* mutants produce a distinct bimodal peak of small (LDL-like) ABCLs (Fig. 3a-c). This pattern warrants further investigation, but may indicate that these alleles directly produce small lipoproteins from the YSL, which are subsequently lipolyzed to produce the second peak. Further study of this allele may provide insight into the specific functional domains of MTP that regulate the size of nascent ABCLs.

### Organ-level changes in lipoprotein localization

The localization of atherogenic lipoproteins throughout an intact organism has not previously been reported. Several observations were in line with our expectations, in that lipoproteins were clearly visible in lipoprotein-producing tissues such as the liver, intestine, and yolk-syncytial layer. Lipoproteins were also clearly evident throughout the circulatory system as expected. However, one of the most striking patterns of lipoprotein localization is the chevron pattern outlining the somites in the trunk region. We suspect this pattern may correspond to the myosepta [45], which include tendinous structures connecting the body segments. Lipoproteins have previously been shown to accumulate in tendons in cases of severe hyperlipidemia [46], but these images suggest that an unexpectedly large fraction of ABCLs localize to these structures in a normal physiological state. The physiological consequences of this association are still unknown, but suggest that the pool of non-circulating ABCLs cannot be ignored in studies of whole-body energy homeostasis.

In addition to this somite pattern, surprisingly strong LipoGlo signal was also observed in the central nervous system (both the brain and spinal cord), colocalizing with the CNS marker *Tg(Xla.Tubb2:mapple-CAAX*). While this pattern was evident throughout development, it was most pronounced in larvae that have low levels of atherogenic lipoproteins throughout the body (such as wild-type larvae at 6 dpf, or larvae that have been fasted or treated with lomitapide for 48 hours, Fig. 5d-f). Further studies will be required to determine the precise brain regions and structures that interact with ABCLs, but the localization pattern observed is strikingly similar to that of fluorescein (a fluorescent dye) after it is injected into the larval zebrafish ventricle [47]. Previous work in mammals has shown that although the blood-brain barrier expresses the LDL-receptor, levels of ApoB within the cerebrospinal fluid are extremely low [48]. These observations suggest that ABCLs may be preferentially retained in the ventricle as a protected source of lipid for the brain in states of nutrient scarcity.

Although atherogenic lipoproteins are most often studied in the bloodstream as a means of evaluating cardiovascular disease risk, these particles interact with essentially every tissue in the body and are involved in numerous processes such as development [49], vision [50], angiogenesis [32], heart function [51], hematopoiesis [52], infection and immunity [53, 54], cancer [55], and diabetes [56]. LipoGlo is an essential tool for broadening the scope of atherogenic lipoprotein biology beyond the current focus on circulating particles. Although the vast majority of NanoLuc protein appears to be associated with intact ABCLs, we cannot eliminate the possibility that some of the protein is cleaved from the particle and may localize independently of ABCLs. Thus these preliminary observations warrant further investigation and validation with orthogonal techniques.

### Characterization of lipoprotein abnormalities in *pla2g12b^-/-^* mutants

Phospholipase A2 group XII B (Pla2g12b) is a catalytically inactive member of the phospholipase gene family. Although this gene lacks catalytic activity and has no other known function, its high level of evolutionary conservation suggests it may have evolved a new function. Previous studies in mice have shown that disruption of *pla2g12b* results in decreased secretion of hepatic triglyceride and ApoB [57], as well as reduced levels of HDL-cholesterol [58], indicating that this gene may play a role in lipoprotein secretion. LipoGlo revealed that *pla2g12b^-/-^* mutant larvae exhibited significantly lower levels of ApoB at multiple stages, and show enrichment of visceral ApoB-NanoLuc levels, both of which are consistent with previously reported defects in lipoprotein secretion. However, evaluation of the lipoprotein size distribution in *pla2g12b^-/-^* mutants revealed bias towards production of small LDL-like particles, which has not been reported previously. Further investigation will be required to determine whether smaller ABCLs are produced directly by the YSL/liver, or if the decreased ABCL secretion rate results in rapid lipolysis of the few lipoproteins that are successfully secreted. As Pla2g12b modulates both lipoprotein size and number, variants in this gene may modulate risk for cardiovascular disease. Ongoing work is exploring both the mechanism of action of this poorly understood protein, as well as the greater physiological repercussions of this mutation.

### Overview of present and future application of LipoGlo

Overall, this study provides several key insights. Firstly, covalent tags to ApoB enable highly sensitive and specific monitoring of ABCLs without disrupting lipoprotein homeostasis. Secondly, the larval zebrafish represents a powerful new model to study ABCLs, as developmental stages provide a highly reproducible pattern of nutrient availability that shows human-like responses to genetic, dietary, and pharmacological stimuli. Additionally, larval zebrafish have unparalleled tractability for genetic and small-molecule screening as well as whole-organism imaging, facilitating the application of these powerful discovery techniques to the field of lipoprotein biology. ABCLs also show significant patterns of enrichment outside of the circulatory system (in association with the somite junctions and central nervous system), laying the foundation for the continued study of extravascular ABCLs. Lastly, *pla2g12b* is a highly conserved regulator of lipoprotein biogenesis that plays a central role in the regulation of the lipoprotein size/subclass distribution. The LipoGlo system thus represents an essential tool to expand our understanding of atherogenic lipoproteins and accelerate the discovery of new approaches to combat atherosclerotic cardiovascular disease.

## ACKNOWLEDGEMENTS

We would like to thank Promega Corp. for providing the NanoLuc plasmid, as well as sample reagents and technical advice that were essential to assay development, as well as serving as a co-sponsor for a local lipid research conference where this work was presented. We would also like to thank Michael Sepanski for collecting the electron micrographs, Dr. Marnie Halpern for providing the unpublished *Tg(Xla.Tubb2:mapple-CAAX*) pan-neuronal marker line and valuable advice on the manuscript, Dr. Yury Miller for providing the *apoC2* mutant line, and the Sanger Institute Zebrafish Mutation project for providing the *pla2g12b* mutant line (sa659). Support was also provided by the National Institutes of Health (R01DK093399 [S.A.F.] and R01DK116079 [S.A.F.]), National Heart, Lung, and Blood Institute (F31HL139338 [J.H.T]) and National Institute of General Medical Sciences (R01GM63904 [S.C.E] & [S.A.F] and P30DK084567 [S.C.E.]). This content is solely the responsibility of the authors and does not necessarily represent the official views of NIH. Additional support for this work was provided by the Carnegie Institution for Science endowment and the G. Harold and Leila Y. Mathers Charitable Foundation (S.A.F).

## AUTHOR CONTRIBUTIONS

J.H.T. and S.A.F. conceived and designed the project, and met frequently to discuss results, plan experiments, and troubleshoot protocols. S.C.E. provided critical reagents and expertise to design and synthesize the TALENs used to create the LipoGlo fish line. J.H.T. executed the experiments, analyzed the results, and wrote the original draft of the paper. J.H.T., S.C.E. and S.A F. revised, edited, and approved the final submitted version of the manuscript.

## MATERIALS AND METHODS

### Zebrafish husbandry and maintenance

Adult zebrafish were maintained on a 14 h light – 10 h dark cycle and fed once daily with ~3.5% body weight of Gemma Micro 500 (Skretting USA). All genotypes were bred into the wild-type AB background. All assays were performed on larvae heterozygous for the ApoB-Nanoluc reporter unless otherwise noted. To monitor the wild-type lipoprotein profile throughout larval development, pairwise crosses were set up between wild-type AB adults and adults homozygous for the ApoB-NanoLuc reporter (apoBb.1^NLuc/NLuc^). To characterize the lipoprotein profile of *mtp* mutant larvae [32], pairwise crosses were set up between mtp^stl/+^ and mtp^stl/+^; apoBb.1^NLuc/NLuc^ adults. To characterize the lipoprotein profile of *apoC2* mutant larvae [15], pairwise crosses were set up between *apoC2^sd38/sd38^* and apoC2^sd38/+^; apoBb.1^NLuc/+^ adults and larvae positive for the NanoLuc reporter were selected for analysis. To characterize the lipoprotein profile of *pla2g12b* mutant larvae [43], pairwise crosses were set up between *pla2g12b^sa659/sa659^* and pla2g12b^sa659/+^; apoBb.1^NLuc/+^ adults. To evaluate association between the ABCLs and the central nervous system, adults homozygous for the ApoB-NanoLuc reporter (apoBb.1^NLuc/NLuc^) were crossed to adults heterozygous for the central nervous system marker *Tg(Xla.Tubb2:mapple-CAAX*), and embryos were screened for mApple prior to fixation and mounting (unpublished reagent provided by the Halpern Lab, c583). As zebrafish sex cannot be determined during the larval stages, gender can be excluded as a variable. All procedures were approved by the Carnegie Institution Animal Care and Use Committee (Protocol #139).

### Genome editing

Genome integration was achieved by co-injection of 500 pg of TALEN mRNA and 30 pg of donor plasmid into 1-cell stage embryos (Supplementary Fig. 2a). Two pairs of TALENs were designed and cloned that target a BsrI restriction site just upstream of the endogenous stop codon of ApoBb.1 using the Mojo Hand design tool [59] and FusX assembly system [60]. TALENs were *in-vitro* transcribed using the T3 mMessage mMachine kti (ThermoFisher Scientific, AM1348) and injected into 1-cell stage zebrafish embryos. Cutting efficiency was quantified by monitoring the loss of BsrI digestion as a result of TALEN nuclease activity, and found to be significantly higher in TALEN pair 2, so this pair was used for genome integration efforts (Supplementary Fig. 2b). A donor plasmid was cloned using 3-fragment MultiSite gateway assembly (Invitrogen, 12537-023) with a 5’ entry element of ~500 bp of the genomic sequence upstream of the ApoBb.1 stop codon, a middle-entry element consisting of in-frame NanoLuc coding sequence, and a 3’ element of ~700 bp of genomic sequence downstream of the ApoBb.1 stop codon [61]. Injected embryos were raised to adulthood and progeny were screened for NanoLuc activity and in-frame fusion of the NanoLuc reporter at the target locus (Supplementary Fig. 2c).

### Preparation and storage of larval homogenate

Individual larvae are homogenized in a standard volume of ABCL stabilization buffer (100 μL). The ABCL stabilization buffer (see recipes) contains cOmplete Mini, EDTA-free Protease Inhibitor Cocktail (Millipore-Sigma, 11836170001), pH buffer and calcium chelator (EGTA, pH 8), and cryoprotectant [62] (Sucrose) to preserve sample integrity during homogenization (Supplementary Fig. 6). The buffer is made as a 2x stock, and larvae are anesthetized in tricaine and placed into tubes in a 50 μL volume and an equal volume of chilled 2x buffer is then added just prior to homogenization. Low-throughput homogenization can be achieved in 1.5 mL centrifuge tubes with disposable pellet-pestles (Fisher scientific, 12-141-363). For high-throughput sample processing, larvae and ABCL stabilization buffer are dispensed into individual wells of a 96-well non-skirted PCR-plate (USAScientific, #1402-9589), sealed with microSeal ‘B’ plate sealing film (Bio-Rad, msb1001), and homogenized in a microplate-horn sonicator (Qsonica, Q700 sonicator with 431MPX microplate horn assembly). For sonication, the plate was placed in the microplate horn filled with 17 mm of chilled RO water and processed at 100% power for a total of 30 seconds, delivered as 2-second pulses interspersed with 1-second pauses. Homogenate was stored on ice for immediate use, or frozen at −20° C and thawed on ice for later use.

### Quantification of ApoB-NanoLuc levels using a plate reader

To quantify ApoB-NanoLuc levels, homogenate (40 μL) was mixed with an equal volume of diluted NanoLuc buffer (for specific dilution see recipes and technical note on NanoLuc buffer) in a 96-well opaque white OptiPlate (Perkin-Elmer, 6005290). Black plates can be used as an alternative that will significantly lower absolute signal intensity, but also reduce light contamination into adjacent wells. The plate was read within 2 minutes of buffer addition using a SpectraMax M5 plate reader (Moleculardevices) set to top-read chemiluminescent detection with a 500 ms integration time. This plate-based assay has a wide linear range and long half-life (Supplementary Fig. 7a-c). However, degree of pigmentation has a significant effect on signal intensity, so this variable should be accounted for with a standard curve or pigment-matched controls should be used as a baseline for comparison (Supplementary Fig. 7d).

### Quantification of lipoprotein size distribution with LipoGlo-electrophoresis

To quantify the electrophoretic mobility of ABCLs, 3% native polyacrylamide gels were cast in Bio-rad mini-protean casting rigs using 1 mm spacer plates and 10-well combs (see recipes). Gels were allowed to polymerize overnight at 4°C and used within 24 h of casting. Each gel included a migration standard comprised of Di-I labeled human LDL (L3482, ThermoFisher Scientific) that was diluted in cryoprotectant and stored in frozen aliquots (see recipes). Gels were assembled into mini-protean electrophoresis rigs at 4°C, filled with pre-chilled 1x TBE and pre-run at 50 V for 30 minutes to equilibrate the gel prior to sample addition. 12 μL of homogenate was then combined with 3 μL of 5x load dye (see recipes), and 12.5 μL of the resulting solution was loaded per well (which corresponds to 10% of the larval homogenate per lane). Gels were then run at 50 V for 30 minutes, followed by 125 volts for 2 h.

Gels were imaged within 1 h of completion of the run. To image each gel, the thin glass short plate was carefully separated from the front of the gel with a gel releaser wedge (see technical note on hydrophobic coating of short plates). With the gel resting on the thick spacer plate, 1 mL of TBE supplemented with 2 μL of Nano-Glo substrate was gently pipetted onto the gel surface. The gel imaging solution was spread evenly across the gel surface with a thin plastic film cut to the size of the spacer plate (Staples, Sliding bar report covers). After a 5-minute equilibration, the gel was placed into an Odyssey Fc (LI-COR Biosciences) gel imaging system (See technical note on gel imaging) and imaged in the chemiluminescence channel for 2 minutes (NanoLuc detection) and then the 600 channel for 30 seconds (Di-l LDL standard detection). Raw images were exported as zip files for further analysis.

The provided gel quantification template (Supplemental File 1) can be used to bin the complex lipoprotein size distribution into biologically relevant groups for analysis, and detailed instructions are provided within the supplemental file. In short, each lane was converted to a plot profile in ImageJ, and divided into LDL, IDL, VLDL, and ZM bins based on migration relative to the Di-I standard, and pixel intensity was summed within each bin for analysis.

### Larvae fixation and imaging

To determine the whole-organism localization of ABCLs, intact larvae are anesthetized and fixed in 4% PFA (diluted in PBS) for 3 h at room temperature. Following fixation, larvae are rinsed 3 times for 15 minutes each in PBS-tween (PBS containing 0.1% tween-20 detergent) and imaged within 12 h of fixation. Agarose for mounting is prepared by melting 0.1 grams of low-melting point agarose (BP160-100, Fisher Scientific) in 10 mLs of 1x TBE. Aliquots are maintained in the liquid state at 42°C in a heat block. Just prior to mounting, agarose aliquots were supplemented with 1% Nano-Glo substrate (furimazine). Fixed larvae are arrayed in droplets on a petri dish lid, and the excess liquid is removed and quickly replaced with a 50 μL droplet of low-melt agarose containing Nano-Glo substrate (1%). The sample is then oriented properly with a flexible poker until the agarose solidifies sufficiently to hold the sample in place. This process was repeated for up to 15 larvae in parallel prior to imaging.

To image the ABCL localization, a Zeiss Axiozoom V16 microscope V16 equipped with a Zeiss AxioCam MRm was set to 30x magnification, 2×2 binning and 2x gain (to increase sensitivity), and programmed to collect a single brightfield exposure (2.4 ms, 10% light intensity) followed by two chemiluminescent imaging exposures (10 and 30 seconds, respectively) with no illumination to collect the NanoLuc signal (See technical note on NanoLuc imaging). Images were quantified in ImageJ by using the brightfield exposure to draw regions of interest (viscera, trunk, and head) and calculating the NanoLuc intensity within each of those ROIs for 30 second chemiluminescent exposure, unless saturated pixels were detected in which case the 10 second exposure was used.

### Density-gradient ultracentrifugation

A density gradient ultracentrifugation (DGUC) protocol was developed by adapting previously published protocols using a 3-layer iodixanol gradient to function with smaller volumes of input sample [40]. Individual larvae were sonicated in 100 μL of sucrose-free ABCL buffer (see recipes) to avoid disruption of the density gradient with sucrose. 15 larvae were pooled per experiment into a single 1.5 mL centrifuge tube and centrifuged for 5 minutes at 6,000 rcf to remove large cellular debris. 1 mL of the resulting supernatant was transferred to a separate tube containing 500 μL of Optiprep Density gradient medium (D1556, Sigma-Aldrich) to yield a 20% iodixanol solution. A 9% iodizanol solution was prepared by adding 1.5 mL of Optiprep to a 15 mL conical tube containing 8.5 mL HEPES-buffered saline (HBS, see recipes), and a 12% solution was prepared by mixing 2 mL Optiprep with 8 mL HBS. A 4.9 mL Optiseal tube (formerly polyallomer, 362185, Beckman-Coulter) was then loaded with 1.5 mL of 9% iodixanol/HBS solution. This solution was carefully underlayered with 1.5 mL of the 12% iodixanol solution using a p1000 pipette fit with both the appropriate p1000 tip as well as a tapered gel loading tip which functioned as a disposable plastic cannula (USA Scientific, 1252-0610). Finally, these solutions were underlayered with 1.5 mL of the 20% iodixanol solution containing the zebrafish homogenate. The tube was then topped up with HBS (~500 uL) so that no air remained and sealed with a cap. Balanced tubes were then loaded into a VTi65.2 rotor and centrifuged at 60,000 rpm in a prechilled Beckman Optima XL 80K Ultracentrifuge set to 4°C with maximum acceleration and deceleration rates.

Following ultracentrifugation, density fractions were collected by carefully piercing the bottom of the tube with a thumbtack, and drip-eluting the samples into 10 separate fractions of approximately 500 μL each. The refractive index of each fraction was determined using a Bausch and Lomb refractometer, and used to calculate solution density using the formula density = 3.3508 x (refractive index) - 3.4675. Fractions were stored on ice or at 10°C, and used within 24 h for a plate-based NanoLuc assay, LipoGlo-electrophoresis, and negative-staining electron microscopy. Note that the high protein and iodixanol content of fraction 1 (highest density) introduces artifacts in the native gel and was therefore excluded, which allowed lane 1 to be dedicated to the Di-I LDL standard.

### Negative-staining electron microscopy

Fractions 4, 7, and 10 from the DGUC experiments outlined above were subjected to negative-staining electron microscopy [41]. 300-mesh copper grids coated with 10 nm formvar and 1 nm carbon (Electron Microscopy Sciences, FCF300-Cu) were ionized using the glow discharge filament in a Denton Vacuum dv-502 evaporator at 75 mTorr for 30 seconds. Anti-capillary forceps were then used to hold the grids in a humidified chamber, and 3 μL of the sample was carefully placed on the surface of the grid and incubated at room temperature for 10 minutes to allow the lipoproteins to adhere to the grid. The grid was then rinsed in 5 droplets of RO-water and then finally 2 droplets of 2% uranyl acetate, and touched lightly to a piece of filter paper to remove excess stain. Grids were imaged at 26,000x magnification on a Tecnai 12 transmission electron microscope.

### DNA extraction and Genotyping

Sonication of zebrafish larvae is a convenient method for highly-parallelized homogenization, as a full plate (96 samples) can be processed simultaneously. However, this process shears DNA into significantly smaller fragments, meaning longer amplicons will amplify less efficiently or not at all. To circumvent this issue, genotyping protocols for this study were designed to use small amplicons (less than 350 bp). If intact DNA is needed for downstream applications, the pellet-pestle method can be used interchangeably with sonication.

DNA extraction of larval homogenate can be achieved with a modified version of the HotShot DNA extraction protocol [63]. 10 μL homogenate is transferred to a pcr tube/plate containing 10 μL of 100 mM NaOH, and heated at 95°C for 20 minutes. The solution was then neutralized with an equal volume (20 μL) of 100 mM Tris pH 8, and either stored frozen (−20 °C)or used immediately as a template for genotyping PCR (2 μL per reaction).

Genotyping was carried out using gene-specific primers (Supplementary Table 1). The ApoBb.1-NanoLuc locus was genotyped using 3 primers with final concentrations as follows: 1 μM primer 9, .2μM primer 10, and .8μM primer 11. This ratio provides similar band intensity for the 113 bp product indicating presence of the WT allele, and the 161 bp product indicating NanoLuc fusion allele (T_a_ = 57°C, extension time 20”) in heterozygotes (only one band will amplify in homozygotes). The *mtp* genotyping locus was amplified using primers 12 and 13 (.5μM each, T_a_ = 60°C, extension time 30”), and digested with 3 units of Avail restriction enzyme, which cuts the mutant (*stlI*) allele. Wild-type zebrafish should have a single 157 bp band, homozygous mutants should have a shorter 129 bp band, and heterozygotes should have both bands present (note the 28 bp fragment is not usually detectable). The *apoC2* genotyping locus was amplified using primers 14 and 15 (.5μM each, T_a_ = 57°C, extension time 30”), and digested with 3 units of Btsαl restriction enzyme, which cuts the WT allele but not the sd38 mutant allele. Wild-type zebrafish should have 102 and 45 bp bands, homozygous mutants should have a single 147 bp band, and heterozygotes should have all 3 bands present. The *pla2g12b* genotyping locus was amplified using primers 16 and 17 (.5μM each, T_a_ = 57°C, extension time 30”), and digested with 3 units of BtsαI restriction enzyme, which cuts the mutant (sa659) allele. Wild-type zebrafish should have a single 150 bp band, homozygous mutants should have a shorter 111 bp band, and heterozygotes should have both bands present (note the 39 bp fragment is not usually detectable).

### Technical notes and troubleshooting

#### NanoLuc Buffer

The NanoLuc enzyme is active in various buffers, but the key consideration is to ensure that substrate is in excess. Manufacturer’s instructions dictate that 1 mL of Buffer plus 20 μL of substrate solution constitutes a 2x buffer, but we have found that this 2x buffer can be diluted 4-fold in PBS and the substrate remains in significant excess.

#### Hydrophobic coating of LipoGlo-electrophoresis plates

The most likely source of artifacts in the <u>LipoGlo-electrophoresis</u> protocol are from stretching or distortion of the fragile 3% polyacrylamide gel while removing the short plate from the gel. To circumvent this issue, the short plates were coated on both sides with Rain-X original glass water repellent (Rain-X, 3.5 oz. bottle). This hydrophobic coating greatly facilitates removal of the short plate while leaving the undistorted gel resting on the spacer plate. This coating is semi-permanent, so it is recommended that a set of coated short plates be dedicated for this purpose and reapplied with coating as needed.

This hydrophobic coating also reduces friction between the short plate and the spacer plate, so it is important that the plates are aligned properly in the casting frames and placed very gently in the casting stands. Too much pressure from the casting stand can cause the plates to slide out of alignment and lead to leaking during casting.

#### Imaging of LipoGlo-electrophoresis gels

The Odyssey Fc offers sensitive signal detection as well as multi-color detection, and is therefore ideal for imaging lipoprotein gels. However, if this equipment is not available, alternative gel imaging systems or a sensitive camera are capable of imaging the gel as well, as the chemiluminescent signal should be detectable by essentially any detector although the exposure time may need to be increased to the order of minutes depending on the sensitivity of the detector. If simultaneous imaging in chemiluminescent and fluorescent channels is not available, a large aliquot of zebrafish homogenate (such as 6 dpf larvae) can be pooled, aliquoted, frozen, and used as an alternative migration normalization standard.

#### NanoLuc imaging

Essentially all background signal in this imaging paradigm comes from two sources: electrical noise from the camera, and light contamination from the environment. Camera noise can be attenuated by using an actively cooled camera and by enabling a blank-subtraction setting to eliminate hot pixels. To reduce contaminating light from the environment, we recommend collecting images in a dark room and shrouding the stage and/or microscope to prevent light from reaching the imaging path. Additionally, we have found that the Zeiss Axiozoom V16 contains infrared emitters and detectors within the imaging path, which result in very high background when long exposures are used. To overcome this issue, we placed a Zeiss BG40 IR blocking filter in front of the camera which effectively filtered the contaminating infrared light.

### Solutions/Recipes

#### ABCL Stabilization Buffer (2x)

For routine preparation of zebrafish homogenate

1 CoMplete mini protease inhibitor tablet
400 μL .5M EGTA (pH8)
1g Sucrose
Adjust volume to 5 mL with reverse osmosis (RO) water

#### Sucrose-free ABCL Stabilization Buffer (2x)

For preparation of zebrafish homogenate for ultracentrifugation

1 CoMplete mini protease inhibitor tablet
400 μL .5M EGTA (pH8)
Adjust volume to 5 mL with RO water

#### Diluted NanoLuc Buffer (2x)

For plate-based measurement of NanoLuc activity

1 mL Nano-Glo buffer
3 mL PBS
20 μL NanoLuc Substrate (furimazine solution)

#### 3% Native Polyacrylamide gels (32 mL, ~4 mini gels)

For <u>LipoGlo-electrophoresis</u> of ABCLs from larval homogenate

22.9 mL RO water
6.4 mL 5x TBE
2.4 mL 40% 19:1 polyacrylamide:bis

→ De-gas under vacuum for 30 minutes
250 μL 10% APS
20 μL TEMED

→ quickly mix by gentle inversion and transfer to casting plates

#### Di-I LDL Lipoprotein migration standard

For normalization of electrophoretic mobility in Ladder Units

200 μL DiI LDL (L3482, Thermofisher Scientific)
4 mL1xTBE
.48 g sucrose (for 10%)
Adjust final volume to 4.8 mL with TBE

→ Divide into 50 μL aliquots and store at −80°C

#### 5x loading dye

For loading homogenate into <u>LipoGlo-electrophoresis</u> gels

4 g sucrose
25 mg bromophenol blue
Adjust to 10 mL with TBE

#### Gel imaging solution (1 gel)

For in-gel chemiluminescent imaging of NanoLuc

1 mLTBE
2 μL furimazine substrate

#### Mounting and Imaging solution (1 mL, ~20 larvae)

For imaging of ABCL distribution in intact larvae

.1g low-melt agarose
10 mL 1x TBE

→ heat in microwave (5-15 seconds) and swirl until dissolved
→ Distribute to 1 mL aliquots in 42°C heat block
Add 10 μL furimazine to 1 mL liquid agarose just prior to mounting

#### HEPES-Buffered Saline

For establishing density gradient for ultracentrifugation

.85g NaCl
10 mL 1M HEPES buffer (pH 7.4)
90 mL RO water

## QUANTIFICATION AND STATISTICAL ANALYSIS

All datasets were initially subjected to Levene’s test for homogeneity of variance. For datasets with a single factor and uniform variance, a one-way ANOVA was used to test for a main effect, and Tukey’s HSD was used for *post hoc* testing. If variance was not uniform (Levene’s <.05), Welch’s ANOVA with a *post hoc* Games-Howell test was used as these tests are robust to the assumption of unequal variance. For two-factor datasets, the Robust Two-Factor ANOVA was used with a *post hoc* Games-Howell test. * denotes p<.01, ** denotes p<.001, and *** denotes p<.0001. For LipoGlo-electrophoresis experiments, statistical tests were run independently for each of the four groups of binned data (ZM, VLDL, IDL, and LDL). In this case, Bonferroni correction was used to adjust for multiple comparisons (corrected significant p<.0125). Bonferroni correction was also applied to the LipoGlo-Microscopy experiments which are binned into three groups, so a significant threshold was set at p<.017. All statistics were run using XLSTAT, with the exception of the Robust Two-Factor ANOVA which was executed in R using the pbad2way function in the WRS2 package (https://cran.r-project.org/web/packages/WRS2/index.html).

## KEY RESOURCES TABLE

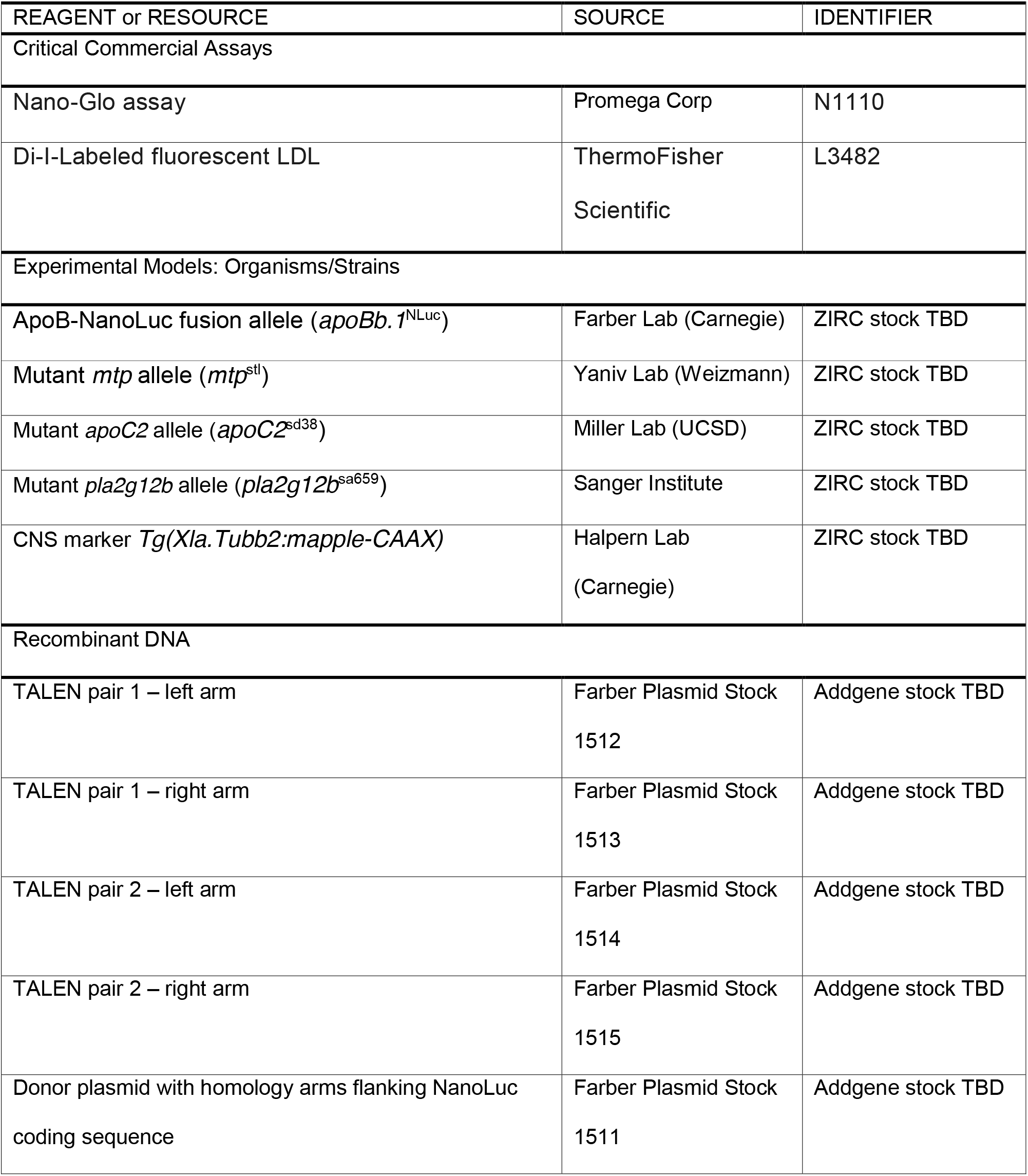

---

**Supplementary Figure 1:**
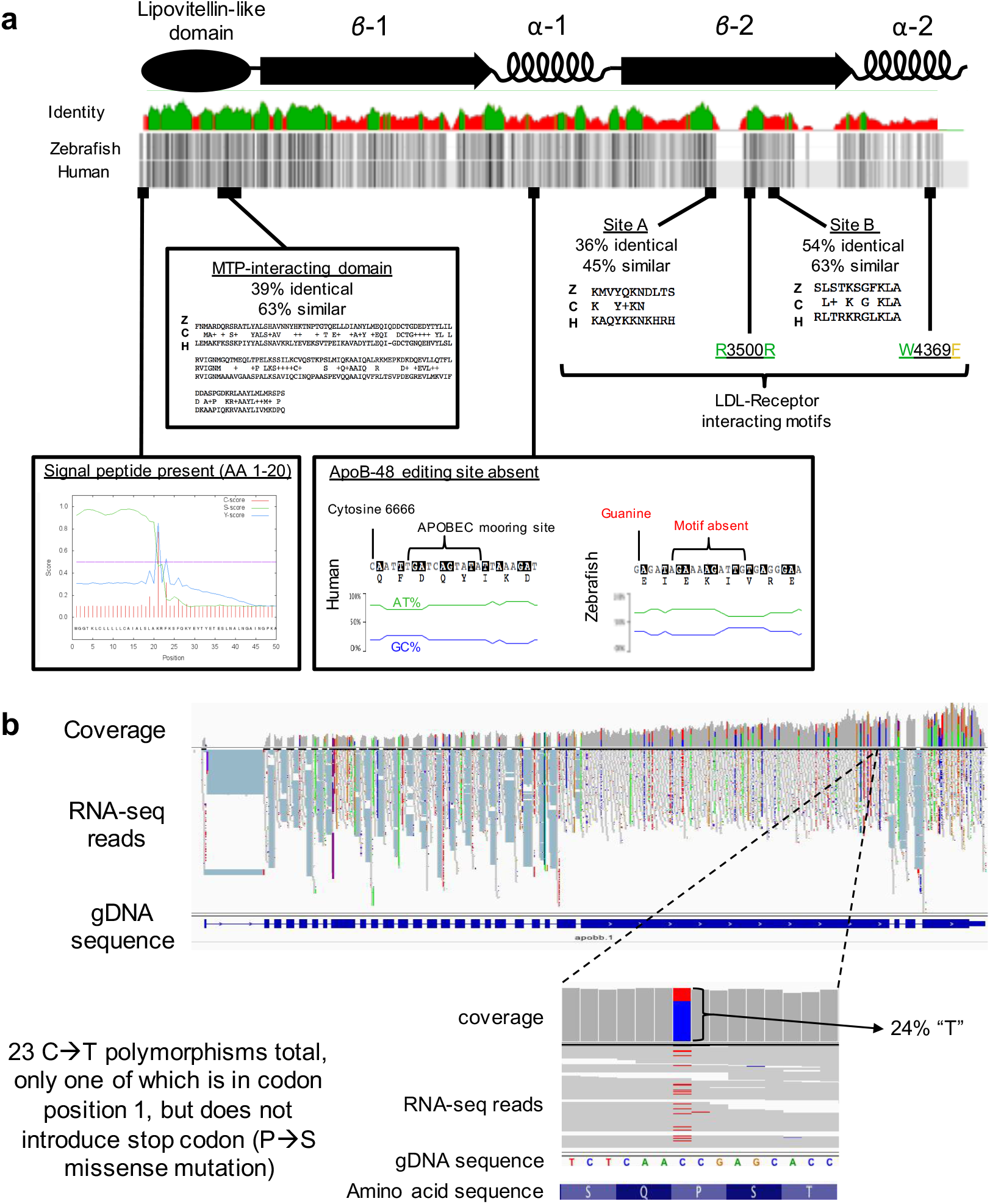
Conservation of functional domains in the zebrafish ApoBb.1 ortholog of Human APOB. **(a)** APOB has a penta-partite domain structure, with an amino-terminal globular domain followed by a series of beta and alpha domains. Consistent with other apolipoprotein sequences, APOB shows relatively low sequence conservation between species at the amino acid level (25% identical, 43% similar, green indicates >30% identity in identity plot). However, sequence conservation is enriched in known ApoB functional domains. For example, there is clear conservation of a signal peptide motif atthe amino terminus. The MTP-interacting domain shows 39% identity and 63% similarity, and the LDL-R interacting motifs are also well-conserved. However, the ApoB-48 editing site appears completely absent, as zebrafish *apoBb.1* lacks the essential C6666 that is edited to form the premature stop, as well as the APOBEC mooring site, and shows only mild AT-richness that has been shown to be important for APOBEC binding **(b)** To further evaluate whether *apoB*-editing takes place in zebrafish, RNA reads were mapped back to this genomic locus. Post-transcriptional C→U editing would appear as a C→T polymorphism in the genomic sequence. 23 instances of C→T polymorphism were observed, but the vast majority (21) appeared in the wobble position (position 3) of the codon as would be expected for true polymorphisms (rather than post-transcriptional RNA-editing). Of the single instance that occurred in position 1, this did not result in a premature stop codon, providing further support for the absence of APOB-editing activity in zebrafish.

**Supplementary Figure 2:**
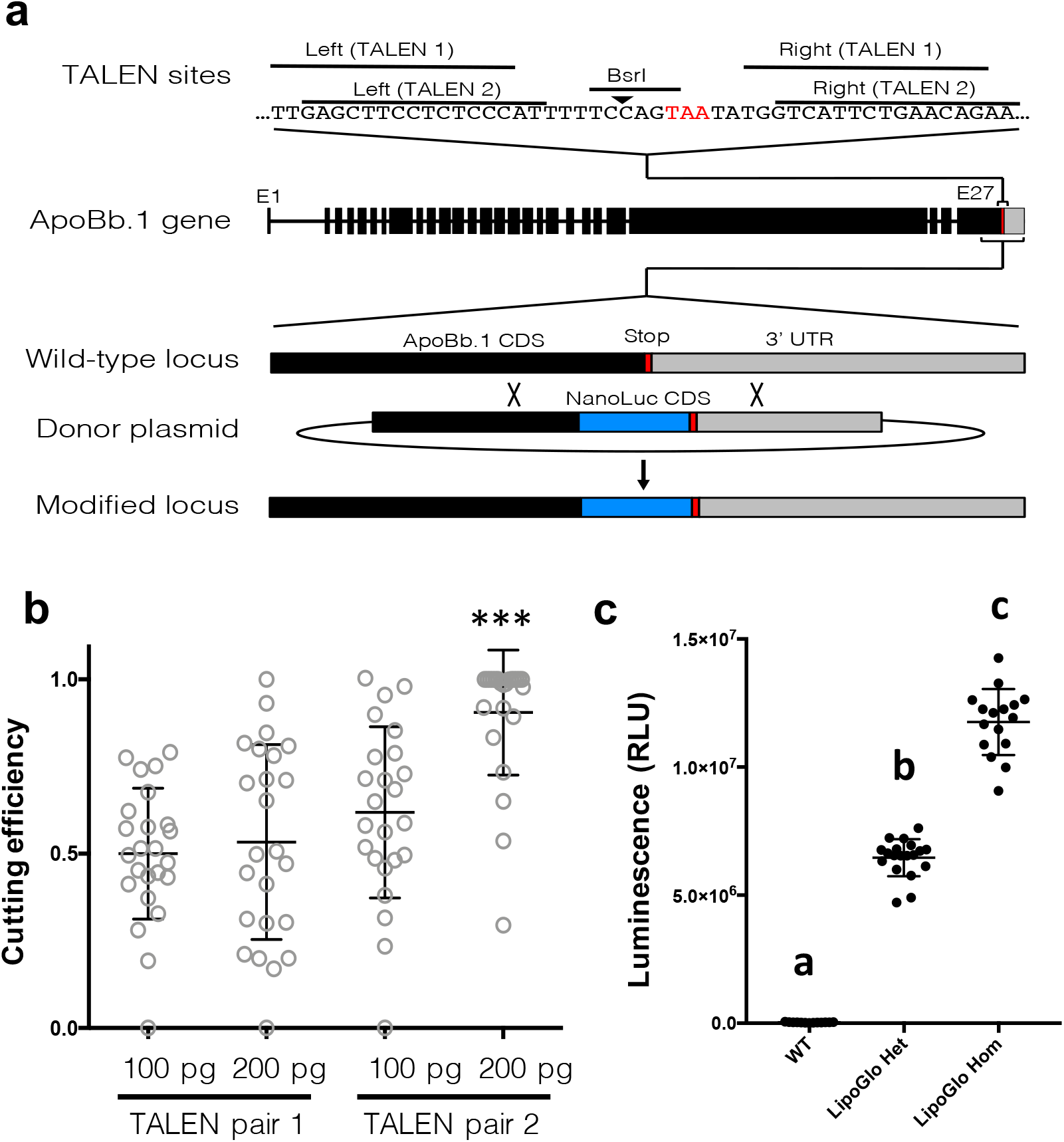
Introduction of an in-frame NanoLuc fusion protein at the endogenous *apoBb.1* locus. **(a)** A BsrI restriction site overlaps partially with the *apoBb.l* stop codon. Two independent pairs of TALENs were designed as shown, and **(b)** tested for cutting efficiency which was quantified as a loss of susceptibility to BsrI digest. TALEN pair 2 showed significantly higher cutting efficiency, and was selected for co-injection with the DNA donor construct (n=24, ANOVA p<0.0001,Tukey’s HSD p<.0001). **(c)** An incross of adult fish heterozygous for the LipoGlo reporter revealed the expected mendelian ratio of offspring, and showed that homozygous carriers produce approximately twice the signal intensity as heterozygotes (12E7±1.3E6 vs 6.5E6±7.3E5) (n≈16, ANOVA p<0.0001,Tukey’s HSD p<.0001). Heterozygous and homozygous carriers of the LipoGlo reporter are viable, fertile, and free of overt morphological defects.

**Supplementary Figure 3:**
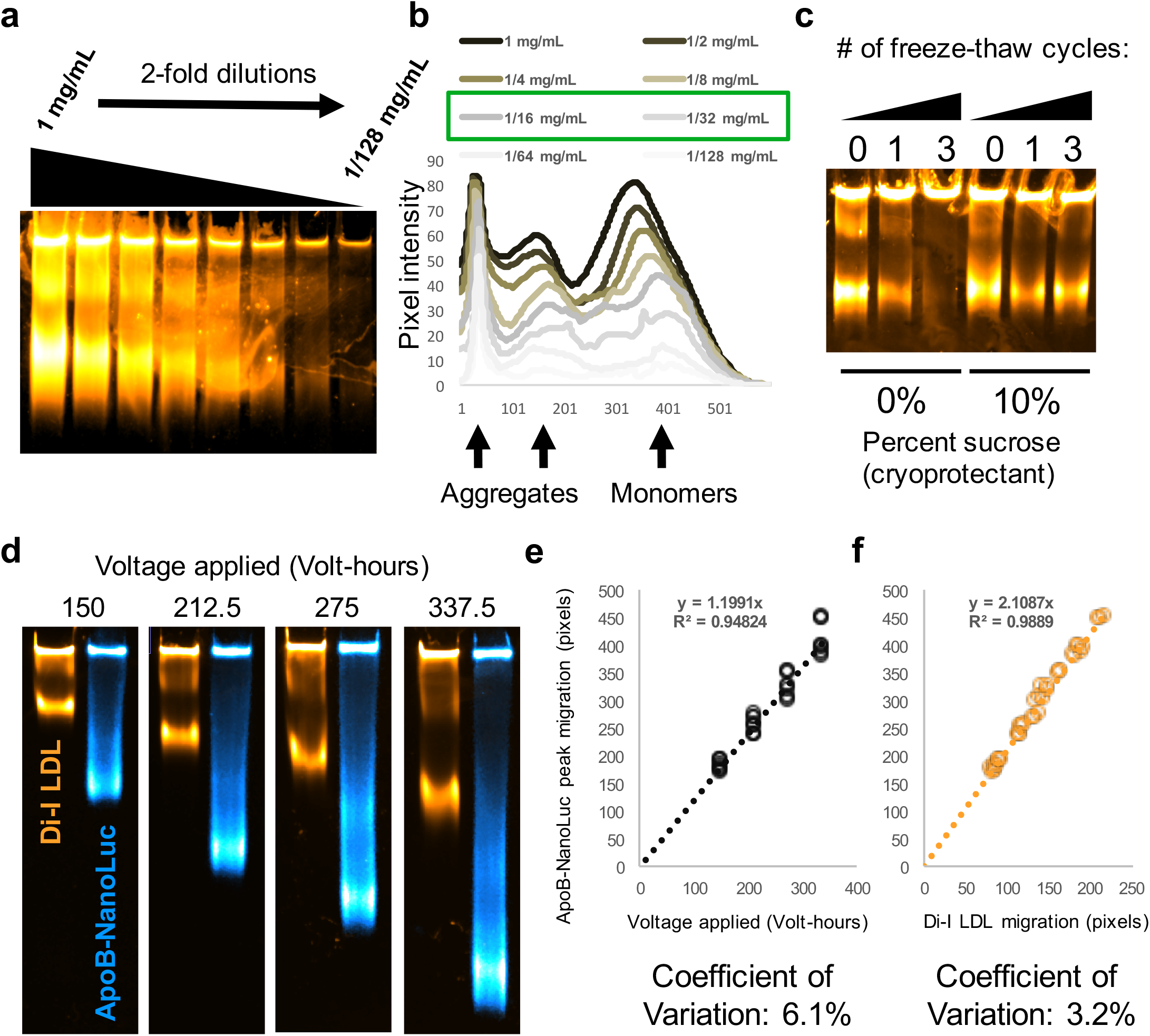
Development of an effective migration standard for lipoprotein gels. It is essential that lipoprotein gels include a ladder or normalization standard that has similar electrophoretic properties to ABCLs. Di-I labeled human LDL serves as a commercially available option that enables standardization not only between multiple gels but also between different labs. **(a)** Di-I LDL was subjected to a series of 2-fold dilutions and separated via Native-PAGE as described and imaged with the Licor-Fc to determine an appropriate dilution factor that was still readily detectable. **(b)** Plot profiles of each of the serial dilutions revealed retardation of peak mobility in the highly concentrated samples, potentially due to overcrowding. Dilution factors between 16 and 32-fold were selected as acceptable (green box), and a 24-fold dilution was used for subsequent assays. **(c)** Sucrose was included as a cryoprotectant during Di-I LDL dilution, and there is no change in peak particle mobility across at least 3 freeze-thaw cycles in the presence of 10% sucrose, whereas the ladder is almost completely aggregated without cryoprotectant. **(d)** To determine the relationship between mobility of the standard and lipoprotein samples, homogenate was prepared and pooled from *mtp* -/- (3 dpf) mutant larvae (which produce primarily LDL-like particles). Samples of homogenate were run alongside Di-I standard for either 150,212.5,275, or 337.5 volt-hours, and the peak migration (in pixels) was quantified for each species. **(e)** While there was a clear linear relationship between ABCL migration and voltage applied (R^2^=.95). **(f)** the relationship was much tighter when electrophoretic mobility was compared to the migration standard (R^2^=.99), validating the utility of Di-I LDL as a migration standard.

**Supplementary Figure 4:**
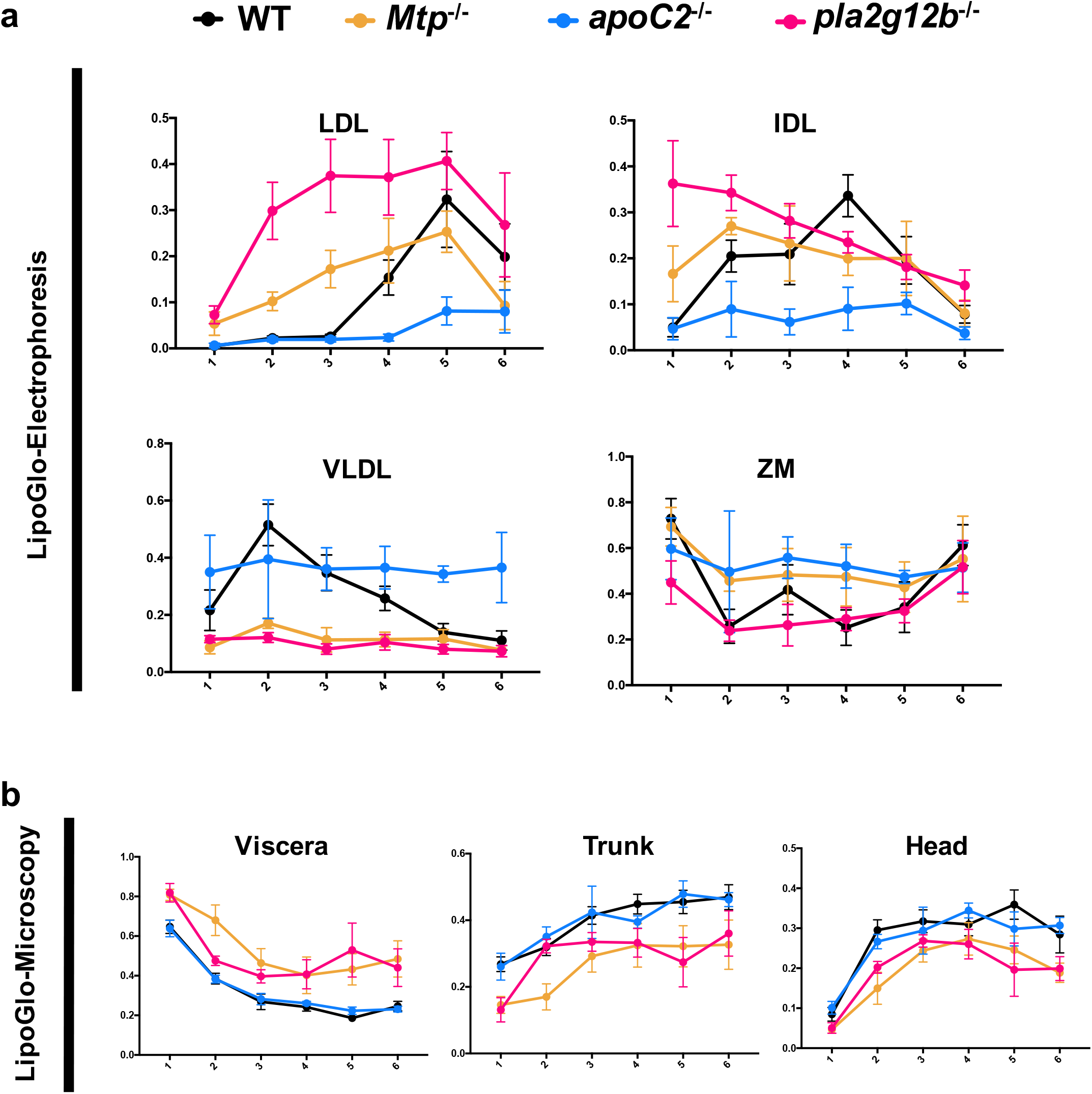
Side-by-side analysis of LipoGlo-Electrophoresis and Microscopy results from mutant genotypes. **(a)** Plots of electrophoresis and **(b)** microscopy data reported in the main text grouped by subclass rather than by genotype and showing standard deviations.

**Supplementary Figure 5:**
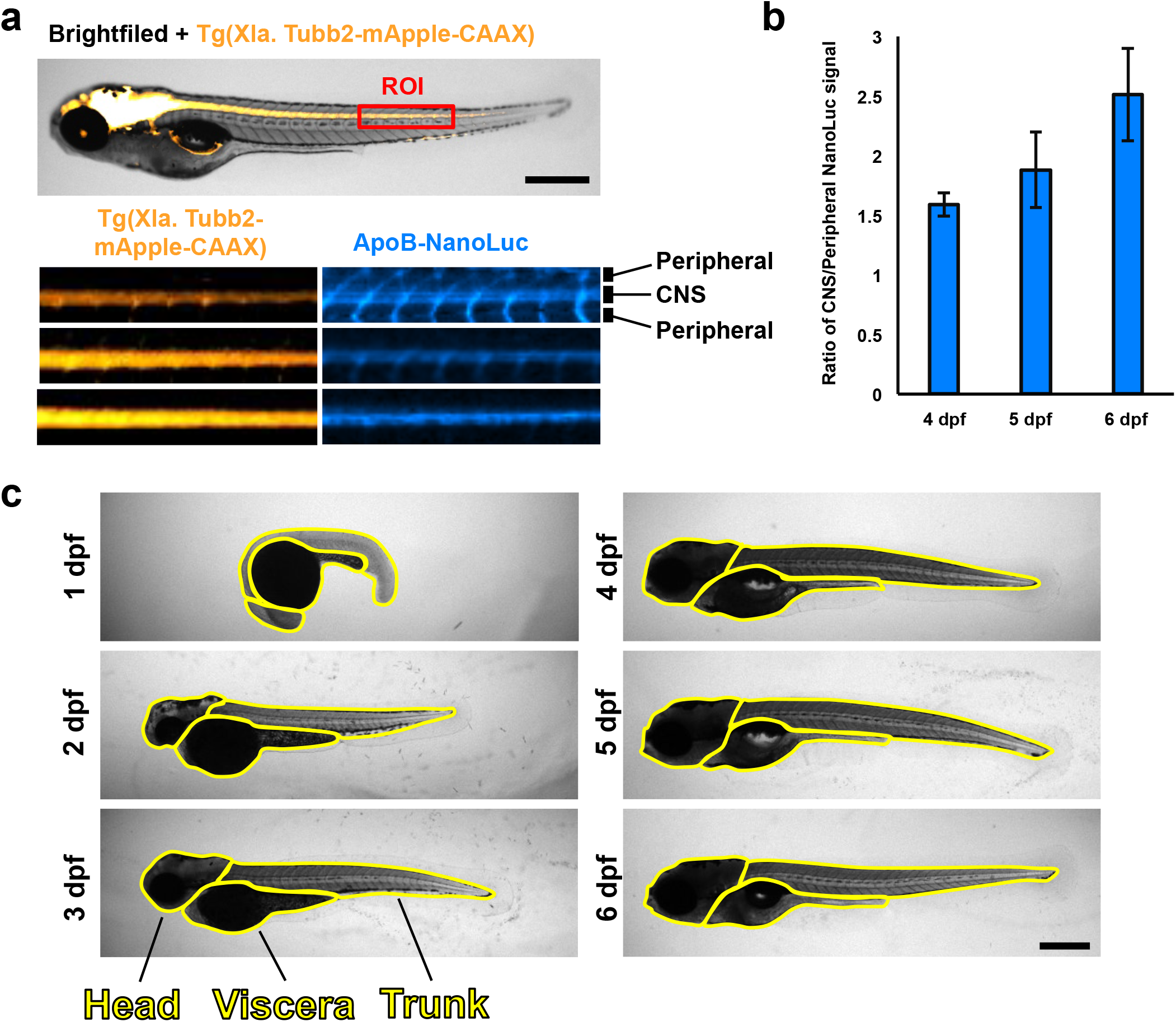
LipoGlo microscopy reveals ABCL localization. **(a)** Three independent clutches of larvae carrying both the CNS marker *Tg(Xla. Tubb2-mApple-CAAX*) and ApoB-NanoLuc fusion were fixed and imaged at 4,5, and 6 dpf as described in the methods section. A 20×100 pixel region of interest (ROI) was drawn centered around the spinal cord (marked by mApple) just distal to the intestine. The mApple and ApoB-NanoLuc channels are displayed separately below (representativeof 15 images pertime point). **(b)** Quantification of the signal intensity in spinal cord (CNS) versus peripheral regions revealed a gradual enrichment of signal in the CNS relative to the periphery from 4-6 dpf (n=15, Welch’s ANOVA p<0.0001, Games-Howell p<.01). **(c)** Representative images of regions of interest corresponding to viscera, trunk, and head regions across development. Scale bars = 500 μm.

**Supplementary Figure 6:**
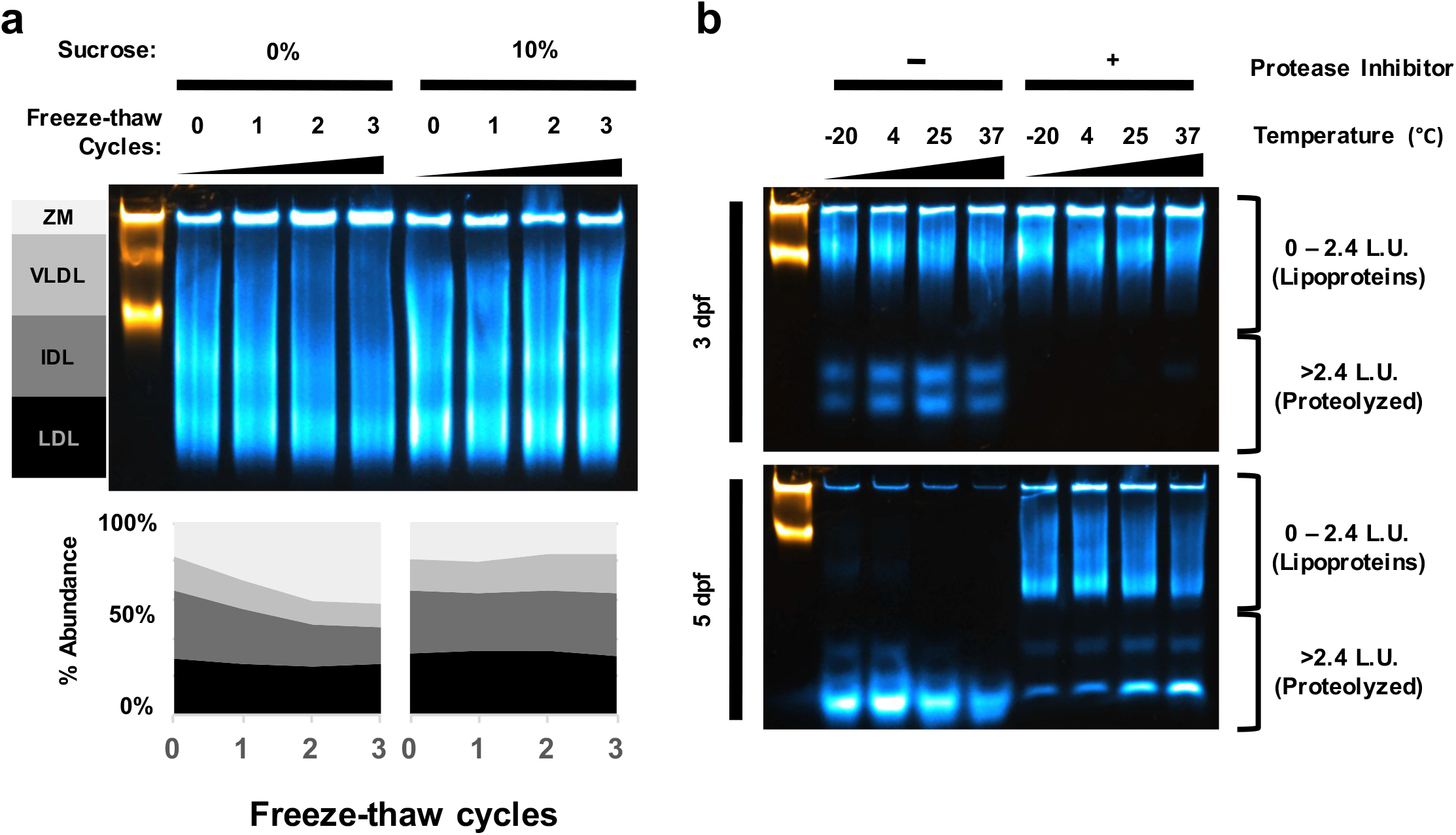
Cryoprotectantand protease-inhibition properties of ABCL stabilization buffer. **(a)** 4 dpf wild-type larvae were homogenized in ABCL stabilization buffer containing 0% or 10% final concentration of sucrose and subjected to between 0 and 3 freeze-thaw cycles, and then separated using <u>LipoGlo-electrophoresis</u> as described in the methods section. While the lipoprotein size distribution remained constant in samples containing sucrose as a cryoprotectant, samples without sucrose showed a gradual enrichment of ZM particles, which appears to be due to aggregation of VLDL and IDL particles. **(b)** Larvae were homogenized in ABCL stabilization buffer with and without the protease inhibitor components (cOmplete mini EDTA-free tablet supplemented with 40 mM final concentration of EGTA, see recipes) and incubated at various temperatures for 2 hours. Samples were then separated by <u>LipoGlo-electrophoresis</u> at 50 V for 30 minutes, and 125 V for 60 minutes. This is 125 Volt-hours less than described in the methods section to enable visualization of proteolysis products. At 3 dpf, protease activity is quite low such that no proteolyzed products are present in the group treated with protease inhibitor, whereas degradation products are visible in a temperature-dependent manner in the absence of inhibitors. By 5 dpf, protease activity is much higher in the homogenate sample, presumably due to development of a mature intestine. Protease activity is still well-controlled in the presence of protease inhibitor at low temperatures, but in the absence of protease inhibitor degradation is so severe that there are signs of both cleavage of NanoLuc from the lipoprotein particle as well as proteolysis of the reporter itself.

**Supplementary Figure 7:**
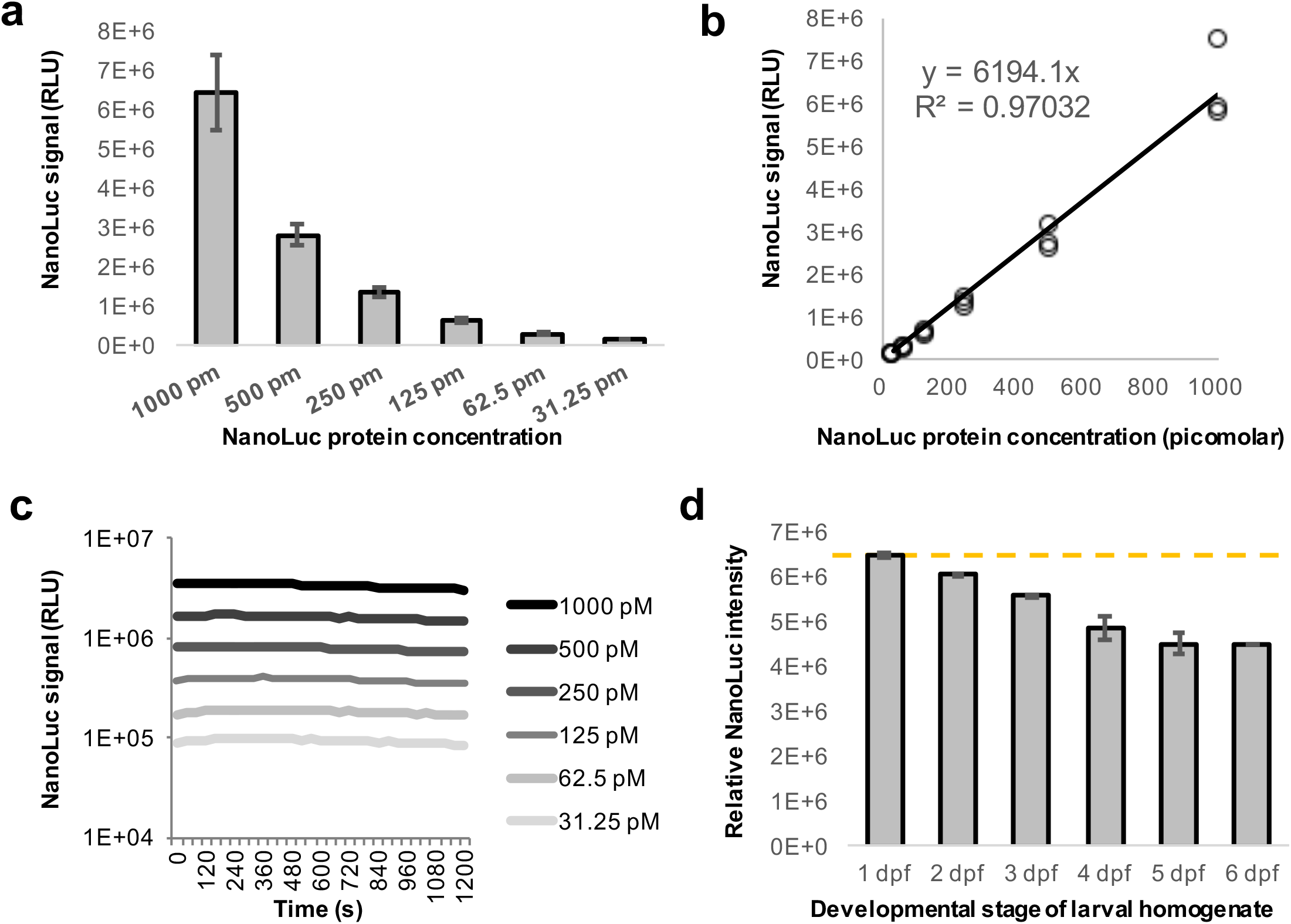
NanoLuc standard curves. **(a)** To determine the absolute concentration of ABCLs in the larval homogenate, purified NanoLuc protein was ordered directly from Promega (NIuc-HT Protein, 5OOug, 54.2KDa, #CSI88401) and diluted to 1 nM working concentration in 1x ABCL stabilization buffer. This solution was subjected to a 6-point series of 2-fold dilutions and used in a plate-based assay for NanoLuc activity, and **(b)** luminescent signal showed excellent linear correlation with protein concentration within this concentration range (R^2^=.97). **(c)** Plate reads were repeated in a kinetic experiment reading well values every 40 seconds for 20 minutes. Signal decayed only marginally in this time window, and half-lives were calculated to be greater than 60 minutes for all concentrations tested. **(d)** There is a marked increase in pigmentation throughout larval development, causing homogenate to become progressively more opaque. To test the effect of pigment on NanoLuc readings, wild-type larvae that lack the ApoB-NanoLuc reporter were homogenized in ABCL stabilization buffer at each day of larval development. This homogenate was then supplemented with a final concentration of 1 nM NanoLuc protein and subjected to a plate read assay. As expected, the relative intensity of NanoLuc signal declines from 1 - 6 dpf, indicating that absolute quantitation of NanoLuc levels should include a standard curve that accounts for changes in larval pigmentation.

**Supplementary Table 1:**
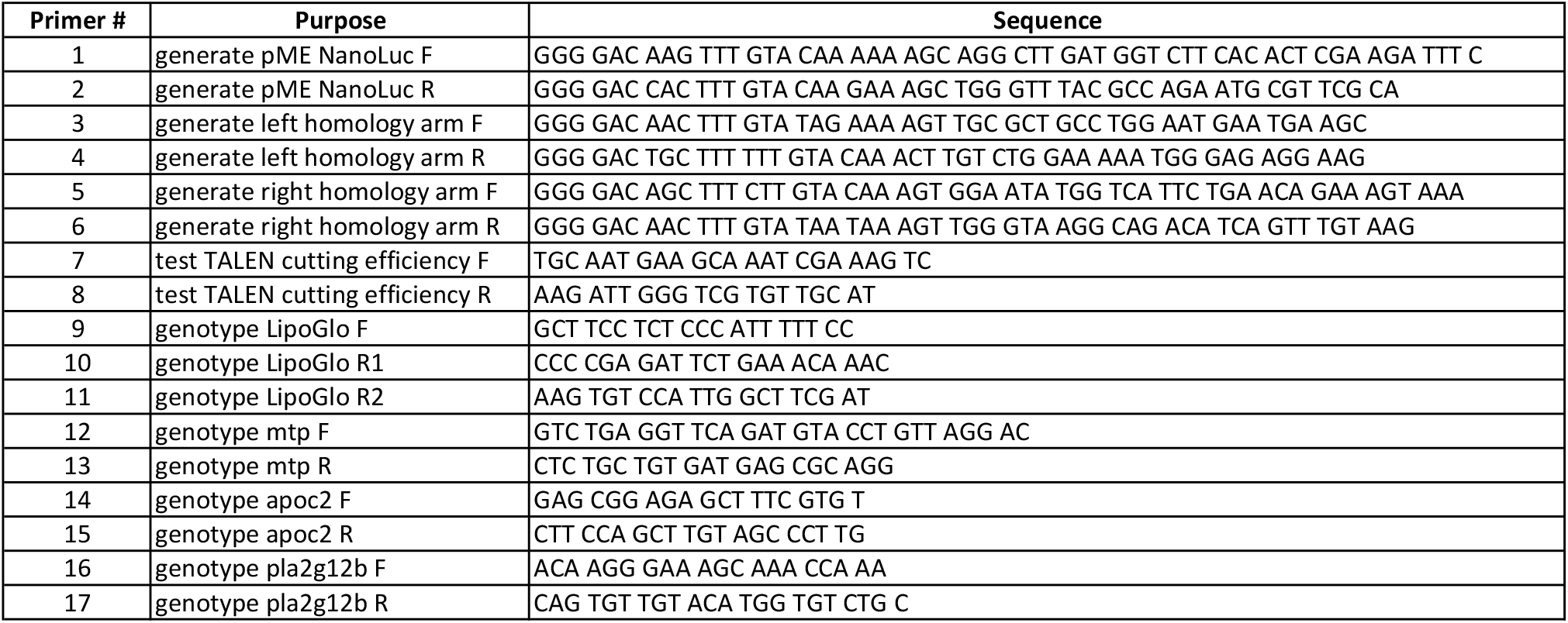
Primers used in this study.

## REFERENCES

1. Tabas, I., K.J. Williams, and J. Boren, Subendothelial lipoprotein retention as the initiating process in atherosclerosis: update and therapeutic implications. Circulation, 2007. 116(16):p. 1832–44.

2. WHO. The top 10 causes of death. Fact Sheets [Web Page] 2018 05/24/2018 [cited 2018 09/25/2018]; Available from: http://www.who.int/news-room/fact-sheets/detail/the-top-10-causes-of-death.

3. Stone, N.J., et al., 2013 ACC/AHA guideline on the treatment of blood cholesterol to reduce atherosclerotic cardiovascular risk in adults: a report of the American College of Cardiology/American Heart Association Task Force on Practice Guidelines. Circulation, 2014. 129(25 Suppl 2): p. S1–45.

4. McQueen, M.J., et al., Lipids, lipoproteins, and apolipoproteins as risk markers of myocardial infarction in 52 countries (the INTERHEART study): a case-control study. Lancet, 2008. 372(9634): p. 224–33.

5. Sniderman, A.D., et al., Discordance analysis of apolipoprotein B and non-high density lipoprotein cholesterol as markers of cardiovascular risk in the INTERHEART study. Atherosclerosis, 2012. 225(2): p. 444–9.

6. Lawler, P.R., et al., Atherogenic Lipoprotein Determinants of Cardiovascular Disease and Residual Risk Among Individuals With Low Low-Density Lipoprotein Cholesterol. J Am Heart Assoc, 2017. 6(7).

7. Rizzo, M. and K. Berneis, Low-density lipoprotein size and cardiovascular risk assessment. QJM, 2006. 99(1): p. 1–14.

8. Austin, M.A., et al., Low-density lipoprotein subclass patterns and risk of myocardial infarction. JAMA, 1988. 260(13): p. 1917–21.

9. Shim, H., et al., A multivariate genome-wide association analysis of 10 LDL subfractions, and their response to statin treatment, in 1868 Caucasians. PLoS One, 2015. 10(4): p. e0120758.

10. Chasman, D.I., et al., Forty-three loci associated with plasma lipoprotein size, concentration, and cholesterol content in genome-wide analysis. PLoS Genet, 2009. 5(11): p. e1000730.

11. Berneis, K.K. and R.M. Krauss, Metabolic origins and clinical significance of LDL heterogeneity. J Lipid Res, 2002. 43(9): p. 1363–79.

12. Rajman, I., et al., LDL particle size: an important drug target? Br J Clin Pharmacol, 1999. 48 (2): p. 125–33.

13. Jacobson, T.A., Opening a new lipid “apo-thecary”: incorporating apolipoproteins as potential risk factors and treatment targets to reduce cardiovascular risk. Mayo Clin Proc, 2011. 86(8): p. 762–80.

14. Xiao, C., et al., Pharmacological Targeting of the Atherogenic Dyslipidemia Complex: The Next Frontier in CVD Prevention Beyond Lowering LDL Cholesterol. Diabetes, 2016. 65(7): p. 1767–78.

15. Liu, C., et al., Apoc2 loss-of-function zebrafish mutant as a genetic model of hyperlipidemia. Dis Model Mech, 2015. 8(8): p. 989–98.

16. Schlegel, A., Zebrafish Models for Dyslipidemia and Atherosclerosis Research. Front Endocrinol (Lausanne), 2016. 7: p. 159.

17. O’Hare, E.A., et al., Disruption of ldlr causes increased LDL-c and vascular lipid accumulation in a zebrafish model of hypercholesterolemia. J Lipid Res, 2014. 55(11): p. 2242–53.

18. Mahley, R.W., Central Nervous System Lipoproteins: ApoE and Regulation of Cholesterol Metabolism. Arterioscler Thromb Vasc Biol, 2016. 36(7): p. 1305–15.

19. Rouault, M., et al., Novel mammalian group XII secreted phospholipase A2 lacking enzymatic activity. Biochemistry, 2003. 42(39): p. 11494–503.

20. Elovson, J., et al., Plasma very low density lipoproteins contain a single molecule of apolipoprotein B. J Lipid Res, 1988. 29(11): p. 1461–73.

21. Fisher, E., E. Lake, and R.S. McLeod, Apolipoprotein B100 quality control and the regulation of hepatic very low density lipoprotein secretion. J Biomed Res, 2014. 28(3): p. 178–93.

22. Kane, J.P., D.A. Hardman, and H.E. Paulus, Heterogeneity of apolipoprotein B: isolation of a new species from human chylomicrons. Proc Natl Acad Sci U S A, 1980. 77(5): p. 2465–9.

23. Davidson, N.O. and G.S. Shelness, APOLIPOPROTEIN B: mRNA editing, lipoprotein assembly, and presecretory degradation. Annu Rev Nutr, 2000. 20: p. 169–93.

24. Otis, J.P., et al., Zebrafish as a model for apolipoprotein biology: comprehensive expression analysis and a role for ApoA-IV in regulating food intake. Dis Model Mech, 2015. 8(3): p. 295–309.

25. Hussain, M.M., et al., Amino acids 430-570 in apolipoprotein B are critical for its binding to microsomal triglyceride transfer protein. J Biol Chem, 1998. 273(40): p. 25612–5.

26. Boren, J., et al., The molecular mechanism for the genetic disorder familial defective apolipoprotein B100. J Biol Chem, 2001. 276(12): p. 9214–8.

27. Hall, M.P., et al., Engineered luciferase reporter from a deep sea shrimp utilizing a novel imidazopyrazinone substrate. ACS Chem Biol, 2012. 7(11): p. 1848–57.

28. Shin, J., J. Chen, and L. Solnica-Krezel, Efficient homologous recombination-mediated genome engineering in zebrafish using TALE nucleases. Development, 2014. 141(19): p. 3807–18.

29. Miyares, R.L., V.B. de Rezende, and S.A. Farber, Zebrafish yolk lipid processing: a tractable tool for the study of vertebrate lipid transport and metabolism. Dis Model Mech, 2014. 7(7): p. 915–27.

30. Hussain, M.M., J. Shi, and P. Dreizen, Microsomal triglyceride transfer protein and its role in apoB-lipoprotein assembly. J Lipid Res, 2003. 44(1): p. 22–32.

31. Jong, M.C., M.H. Hofker, and L.M. Havekes, Role of ApoCs in lipoprotein metabolism: functional differences between ApoC1, ApoC2, and ApoC3. Arterioscler Thromb Vasc Biol, 1999. 19(3): p. 472–84.

32. Avraham-Davidi, I., et al., ApoB-containing lipoproteins regulate angiogenesis by modulating expression of VEGF receptor 1. Nat Med, 2012. 18(6): p. 967–73.

33. Cuchel, M., et al., Inhibition of microsomal triglyceride transfer protein in familial hypercholesterolemia. N Engl J Med, 2007. 356(2): p. 148–56.

34. Carten, J.D., M.K. Bradford, and S.A. Farber, Visualizing digestive organ morphology and function using differential fatty acid metabolism in live zebrafish. Dev Biol, 2011. 360(2): p. 276–85.

35. Feingold, K.R. and C. Grunfeld, Introduction to Lipids and Lipoproteins, in Endotext, L.J. De Groot, et al., Editors. 2000: South Dartmouth (MA).

36. Singh, Y., et al., A rapid 3% polyacrylamide slab gel electrophoresis method for high through put screening of LDL phenotype. Lipids Health Dis, 2008. 7: p. 47.

37. Hoefner, D.M., et al., Development of a rapid, quantitative method for LDL subfractionation with use of the Quantimetrix Lipoprint LDL System. Clin Chem, 2001. 47(2): p. 266–74.

38. Sato, A., et al., Angiotensin II induces the aggregation of native and oxidized low-density lipoprotein. Eur Biophys J, 2018. 47(1): p. 1–9.

39. Tiwari, S. and S.A. Siddiqi, Intracellular trafficking and secretion of VLDL. Arterioscler Thromb Vasc Biol, 2012. 32(5): p. 1079–86.

40. Yee, M.S., et al., Lipoprotein separation in a novel iodixanol density gradient, for composition, density, and phenotype analysis. J Lipid Res, 2008. 49(6): p. 1364–71.

41. Garewal, M., L. Zhang, and G. Ren, Optimized negative-staining protocol for examining lipid-protein interactions by electron microscopy. Methods Mol Biol, 2013. 974: p. 111–8.

42. Westerfield, M., The zebrafish book: a guide for the laboratory use of zebrafish (Danio rerio). 2007: University of Oregon press.

43. Kettleborough, R.N., et al., A systematic genome-wide analysis of zebrafish protein-coding gene function. Nature, 2013. 496(7446): p. 494–7.

44. Chico, T.J., P.W. Ingham, and D.C. Crossman, Modeling cardiovascular disease in the zebrafish. Trends Cardiovasc Med, 2008. 18(4): p. 150–5.

45. Charvet, B., et al., Development of the zebrafish myoseptum with emphasis on the myotendinous junction. Cell Tissue Res, 2011. 346(3): p. 439–49.

46. Tsouli, S.G., et al., Regression of Achilles tendon thickness after statin treatment in patients with familial hypercholesterolemia: an ultrasonographic study. Atherosclerosis, 2009. 205(1): p. 151–5.

47. Henson, H.E., et al., Functional and genetic analysis of choroid plexus development in zebrafish. Front Neurosci, 2014. 8: p. 364.

48. Dehouck, B., et al., A new function for the LDL receptor: transcytosis of LDL across the blood-brain barrier. J Cell Biol, 1997. 138(4): p. 877–89.

49. Neumann, S., et al., Mammalian Wnt3a is released on lipoprotein particles. Traffic, 2009. 10 (3): p. 334–43.

50. Pikuleva, I.A. and C.A. Curcio, Cholesterol in the retina: the best is yet to come. Prog Retin Eye Res, 2014. 41: p. 64–89.

51. Yu, X., et al., Inhibition of cardiac lipoprotein utilization by transgenic overexpression of Angptl4 in the heart. Proc Natl Acad Sci U S A, 2005. 102(5): p. 1767–72.

52. Liu, C., et al., Lipoprotein lipase regulates hematopoietic stem progenitor cell maintenance through DHA supply. Nat Commun, 2018. 9(1): p. 1310.

53. Manifold-Wheeler, B.C., et al., Serum Lipoproteins Are Critical for Pulmonary Innate Defense against Staphylococcus aureus Quorum Sensing. J Immunol, 2016. 196(1): p. 328–35.

54. Bashmakov, Y.K., et al., ApoB-containing lipoproteins promote infectivity of chlamydial species in human hepatoma cell line. World J Hepatol, 2010. 2(2): p. 74–80.

55. Borgquist, S., et al., Apolipoproteins, lipids and risk of cancer. Int J Cancer, 2016. 138(11): p. 2648–56.

56. Ley, S.H., et al., Association of apolipoprotein B with incident type 2 diabetes in an aboriginal Canadian population. Clin Chem, 2010. 56(4): p. 666–70.

57. Guan, M., et al., Hepatocyte nuclear factor-4 alpha regulates liver triglyceride metabolism in part through secreted phospholipase A(2) GXIIB. Hepatology, 2011. 53(2): p. 458–66.

58. Aljakna, A., et al., Pla2g12b and Hpn are genes identified by mouse ENU mutagenesis that affect HDL cholesterol. PLoS One, 2012. 7(8): p. e43139.

59. Neff, K.L., et al., Mojo Hand, a TALEN design tool for genome editing applications. BMC Bioinformatics, 2013. 14: p. 1.

60. Ma, A.C., et al., FusX: A Rapid One-Step Transcription Activator-Like Effector Assembly System for Genome Science. Hum Gene Ther, 2016. 27(6): p. 451–63.

61. Petersen, L.K. and R.S. Stowers, A Gateway MultiSite recombination cloning toolkit. PLoS One, 2011. 6(9): p. e24531.

62. Rumsey, S.C., et al., Cryopreservation with sucrose maintains normal physical and biological properties of human plasma low density lipoproteins. J Lipid Res, 1992. 33(10): p. 1551–61.

63. Meeker, N.D., et al., Method for isolation of PCR-ready genomic DNA from zebrafish tissues. Biotechniques, 2007. 43(5): p. 610, 612, 614.

